# Invariant neural subspaces maintained by feedback modulation

**DOI:** 10.1101/2021.10.29.466453

**Authors:** Laura Bella Naumann, Joram Keijser, Henning Sprekeler

## Abstract

Sensory systems reliably process incoming stimuli in spite of changes in context. Most recent models accredit this context invariance to an extraction of increasingly complex sensory features in hierarchical feedforward networks. Here, we study how context-invariant representations can be established by feedback rather than feedforward processing. We show that feedforward neural networks modulated by feedback can dynamically generate invariant sensory representations. The required feedback can be implemented as a slow and spatially diffuse gain modulation. The invariance is not present on the level of individual neurons, but emerges only on the population level. Mechanistically, the feedback modulation dynamically reorients the manifold of neural activity and thereby maintains an invariant neural subspace in spite of contextual variations. Our results highlight the importance of population-level analyses for understanding the role of feedback in flexible sensory processing.

## Introduction

In natural environments our senses are exposed to a colourful mix of sensory impressions. Behaviourally relevant stimuli can appear in varying contexts, such as variations in lighting, acoustics, stimulus position or the presence of other stimuli. Different contexts may require different responses to the same stimulus, for example when the behavioural task changes (context dependence). Alternatively, the same response may be required for different stimuli, for example when the sensory context changes (context invariance). Recent advances have elucidated how *context-dependent* processing can be performed by recurrent feedback in neural circuits (Mante et al., 2013; Wang et al., 2018a; Dubreuil et al., 2020). In contrast, the role of feedback mechanisms in context-*invariant* processing is not well understood.

In the classical view, stimuli are hierarchically processed towards a behaviourally relevant percept that is invariant to contextual variations. This is achieved by extracting increasingly complex features in a feedforward network (Kriegeskorte, 2015; Zhuang et al., 2021; Yamins and DiCarlo, 2016). Models of such feedforward networks have been remarkably successful at learning complex perceptual tasks (LeCun et al., 2015), and they account for various features of cortical sensory representations (DiCarlo and Cox, 2007; Kriegeskorte et al., 2008; DiCarlo et al., 2012; Hong et al., 2016; Cichy et al., 2016). Yet, these models neglect feedback pathways, which are abundant in sensory cortex (Felleman and Van Essen, 1991; Markov et al., 2014) and shape sensory processing in critical ways (Gilbert and Li, 2013). Incorporating these feedback loops into models of sensory processing increases their flexibility and robustness (Spoerer et al., 2017; Alamia et al., 2021; Nayebi et al., 2021) and improves their fit to neural data (Kar et al., 2019; Kietzmann et al., 2019; Nayebi et al., 2021). At the neuronal level, feedback is thought to modulate rather than drive local responses (Sherman and Guillery, 1998), for instance depending on behavioral context (Niell and Stryker, 2010; Vinck et al., 2015; Kuchibhotla et al., 2017; Dipoppa et al., 2018).

Here, we investigate the hypothesis that feedback modulation provides a neural mechanism for context-invariant perception. To this end, we trained a feedback-modulated network model to perform a context-invariant perceptual task and studied the resulting neural mechanisms. We show that the feedback modulation does not need to be temporally or spatially precise and can be realised by feedback-driven gain modulation in rate-based networks of excitatory and inhibitory neurons. To solve the task, the feedback loop dynamically maintains an invariant subspace in the population representation (Hong et al.,2016). This invariance is not present at the single neuron level. Finally, we find that the feedback conveys a nonlinear representation of the context itself, which can be hard to discern by linear decoding methods.

These findings corroborate that feedback-driven gain modulation of feedforward networks enables context-invariant sensory processing. The underlying mechanism links single neuron modulation with its function at the population level, highlighting the importance of population-level analyses.

## Results

As a simple instance of a context-invariant task, we considered a dynamic version of the blind source separation problem. The task is to recover unknown sensory sources, such as voices at a cocktail party (McDermott, 2009), from sensory stimuli that are an unknown mixture of the sources. In contrast to the classical blind source separation problem, the mixture can change in time, for example, when the speakers move around, thus providing a time-varying sensory context. Because the task requires a dynamic inference of the context, it cannot be solved by feedforward networks (Supp. Fig. S1) or standard blind source separation algorithms (e.g., independent component analysis; Bell and Sejnowski, 1995; Hyvärinen and Oja, 2000). We hypothesised that this dynamic task can be solved by a feedforward network that is subject to modulation from a feedback signal. In our model the feedback signal is provided by a modulatory system that receives both the sensory stimuli and the network output (Fig. 1a).

**Figure 1.**
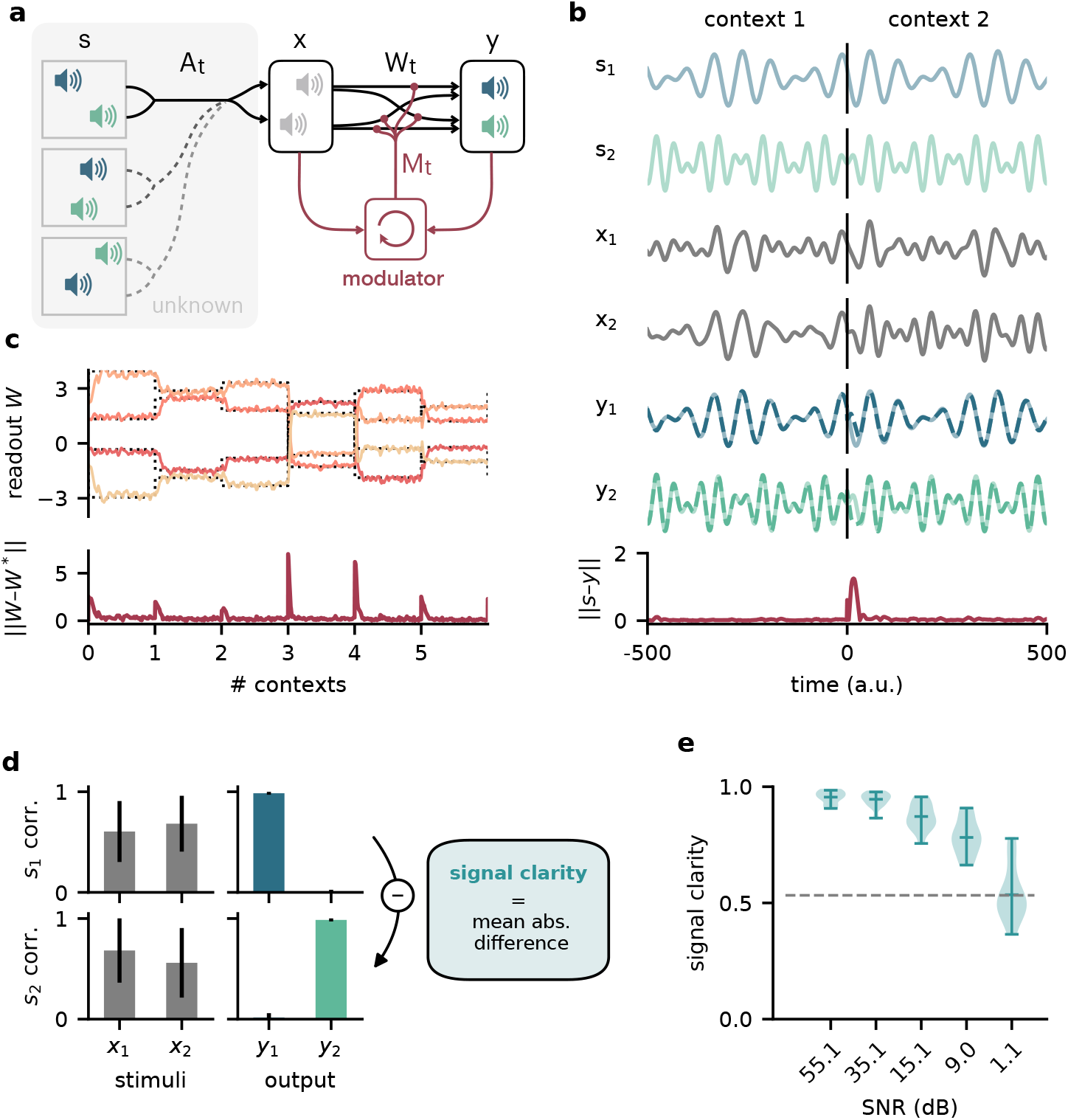
Dynamic blind source separation by modulation of feedforward connections. **a.** Schematic of the feedforward network model receiving feedback modulation from a modulator (a recurrent network). **b.** Top: Sources (*s*_1,2_), sensory stimuli (*x*_1,2_) and network output (*y*_1,2_) for two different source locations (contexts). Bottom: Deviation of output from the sources. **c.** Top: Modulated readout weights across 6 contexts (source locations); dotted lines indicate the true weights of the inverted mixing matrix. Bottom: Deviation of readout from target weights. **d.** Correlation between the sources and the sensory stimuli (left), the network outputs (center), and calculation of the *signal clarity* (right). Errorbars indicate standard deviation across 20 contexts. **e.** Violin plot of the signal clarity for different noise levels in the sensory stimuli across 20 different contexts.

### Dynamic blind source separation by modulation of feedforward weights

Before we gradually take this to the neural level, we illustrate the proposed mechanism in a simple example, in which the modulatory system provides a time-varying multiplicative modulation of a linear two-layer network (see Methods and Models). For illustration, we used compositions of sines with different frequencies as source signals (*s*, Fig. 1b, top). These sources were linearly mixed to generate the sensory stimuli (*x*) that the network received as input; *x* = *A_t_ s* (Fig. 1a, b). The linear mixture (*A_t_*) changed over time, akin to varying the location of sound sources in a room (Fig. 1a). These locations provided a time-varying sensory context that changed on a slower timescale than the sources themselves. The feedforward network had to recover the sources from the mixed sensory stimuli. To achieve this, we trained the modulator to dynamically adjust the weights of the feedforward network (*W*_0_) such that the network output (*y*) matches the sources:

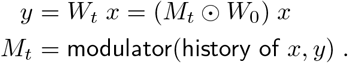

Because the modulation requires a dynamic inference of the context, the modulator is a recurrent neural network. The modulator was trained using supervised learning. Afterwards, its weights were fixed and it no longer had access to the target sources (see Methods & Models, Fig. 8). The modulator therefore had to use its recurrent dynamics to determine the appropriate modulatory feedback for the time-varying context, based on the sensory stimuli and the network output. Put differently, the modulator had to learn an internal model of the sensory data and the contexts, and use it to establish the desired context invariance in the output.

After learning, the modulated network disentangled the sources, even when the context changed (Fig. 1b, Supp. Fig. S2a,b). Context changes produced a transient error in the network’s output, but it quickly resumed matching the sources (Fig. 1b, bottom). The transient errors occur, because the modulator needs time to infer the new context from the time-varying inputs, before it can provide the appropriate feedback signal to the feedforward network (Supp. Fig. S6a, cf. Supp. Fig. S1g-i). The modulated feedforward weights inverted the linear mixture of sources by switching on the same timescale (Fig. 1c).

To quantify how well the sources were separated, we measured the correlation coefficient of the outputs with each source over several contexts. Consistent with a clean separation, we found that each of the two outputs strongly correlated with only one of the sources. In contrast, the sensory stimuli showed a positive average correlation for both sources, as expected given the positive linear mixture (Fig. 1d, left). We determined the *signal clarity* as the absolute difference between the correlation with the first compared to the second source, averaged over the two outputs, normalised by the sum of the correlations (Fig. 1d, right; see Methods and Models). The signal clarity thus determines the degree of signal separation, where a value close to 1 indicates a clean separation as in Fig. 1d. Note that the signal clarity of the sensory stimuli is around 0.5 and can be used as a reference.

We next probed the network’s robustness by adding noise to the sensory stimuli. We found that the signal clarity gradually decreased with increasing noise levels, but only degraded to chance performance when the signal-to-noise ratio was close to 1 (1.1 dB, Fig. 1e, Supp. Fig. S2e).

The network performance did not depend on the specific source signals (Supp. Fig. S3) or the number of sources (Supp. Fig. S4), as long as it had seen them during training. Yet, because the network had to learn an internal model of the task, we expected a limited degree of generalisation to new situations. Indeed, the network was able to interpolate between source frequencies seen during training (Supp. Fig. S5), but failed on sources and contexts that were qualitatively different (Supp. Fig. S6b-d). The specific computations performed by the modulator are therefore idiosyncratic to the problem at hand. Hence, we did not investigate the internal dynamics of the modulator in detail, but concentrated on its effect on the feedforward network.

Since feedback-driven modulation enables flexible context-invariant processing in a simple abstract model, we wondered how this mechanism might be implemented at the neural level. For example, how does feedback-driven modulation function when feedback signals are slow and imprecise? And how does the modulation affect population activity? In the following, we will gradually increase the model complexity to account for biological constraints and pinpoint the population-level mechanisms of feedback-mediated invariance.

### Invariance can be established by slow feedback modulation

Among the many modulatory mechanisms, even the faster ones are believed to operate on timescales of hundreds of milliseconds (Bang et al., 2020; Molyneaux and Hasselmo, 2002), raising the question if feedback-driven modulation is sufficiently fast to compensate for dynamic changes in environmental context.

To investigate how the timescale of modulation affects the performance in the dynamic blind source separation task, we trained network models, in which the modulatory feedback had an intrinsic timescale that forced it to be slow. We found that the signal clarity degraded only when this timescale was on the same order of magnitude as the timescale of contextual changes (Fig. 2a). Note that timescales in this model are relative, and could be arbitrarily rescaled. While slower feedback modulation produced a larger initial error (Fig. 2b,c), it also reduced the fluctuations in the readout weights such that they more closely follow the optimal weights (Fig. 2b). This speed-accuracy trade-off explains the lower and more variable signal clarity for slow modulation (Fig. 2a), because the signal clarity was measured over the whole duration of a context and the transient onset error dominated over the reduced fluctuations.

**Figure 2.**
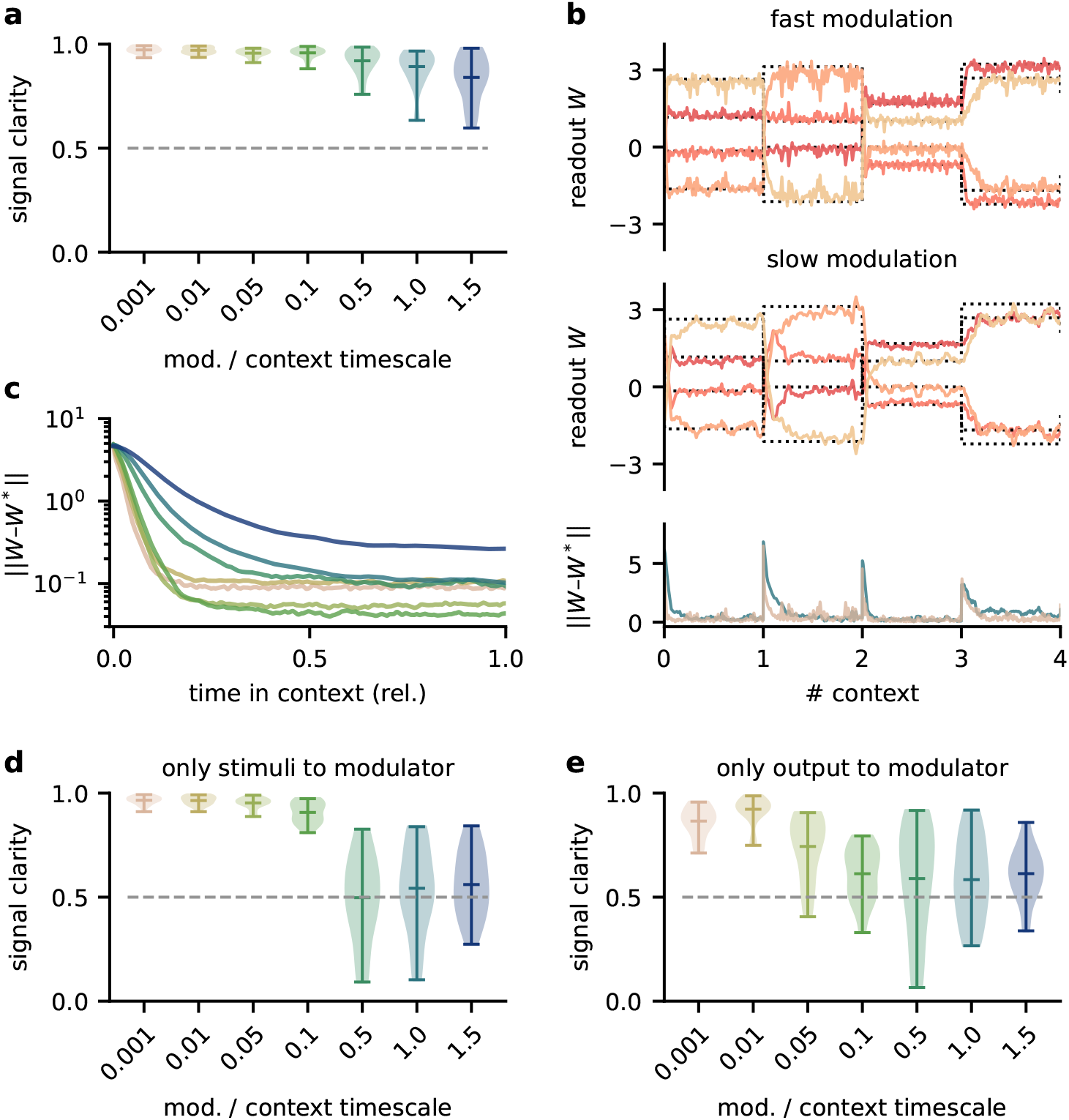
The network model is not sensitive to slow feedback modulation. **a.** Signal clarity in the network output for varying timescales of modulation relative to the intervals at which the source locations change. **b.** Modulated readout weights across 4 source locations (contexts) for fast (top) and slow (center) feedback modulation; dotted lines indicate the optimal weights (the inverse of the mixing matrix). Bottom: deviation of the readout weights from the optimal weights for fast and slow modulation. Colours correspond to the relative timescales in (a). Fast and slow timescales are 0.001 and 1, respectively. **c.** Mean deviation of readout from optimal weights within contexts; averaged over 20 contexts. Colours code for timescale of modulation (see (a)). **d. & e.** Same as (a) but for models in which the modulatory system only received the sensory stimuli *x* or the network output *y*, respectively.

To determine architectural constraints on the modulatory system, we asked how these results depended on the input it received. So far, the modulatory system received the feedforward network’s inputs (the sensory stimuli) and its outputs (the inferred sources, see Fig. 1a), but are both of these necessary to solve the task? We found that when the modulatory system only received the sensory stimuli, the model could still learn the task, though it was more sensitive to slow modulation (Fig. 2d, Supp. Fig. S7). When the modulatory system had to rely on the network output alone, task performance was impaired even for fast modulation (Fig. 2e, Supp. Fig. S7). Thus, while the modulatory system is more robust to slow modulation when it receives the network output, the output is not sufficient to solve the task.

Taken together, these results show that the biological timescale of modulatory mechanisms does not pose a problem for flexible feedback-driven processing, as long as the feedback modulation changes on a faster timescale than variations in the context. In fact, slow modulation can increase processing accuracy by averaging out fluctuations in the feedback signal. Nevertheless, slow modulation likely requires the modulatory system to receive both the input and output of the sensory system it modulates.

### Invariance can be established by spatially diffuse feedback modulation

Neuromodulators are classically believed to diffusely affect large areas of the brain. Furthermore, signals in the brain are processed by populations of neurons. We wondered if the proposed modulation mechanism is consistent with such biological constraints. We therefore extended the network model such that the sensory stimuli are projected to a population of 100 neurons. A fixed linear readout of this population determined the network output. The neurons in the population received spatially diffuse modulatory feedback (Fig. 3a) such that the feedback modulation affected neighbouring neurons similarly. We here assume that all synaptic weights to a neuron receive the same modulation, such that the feedback performs a gain modulation of neural activity (Ferguson and Cardin, 2020). The spatial specificity of the modulation was determined by the number of distinct feedback signals and their spatial spread (Fig. 3b, Supp. Fig. S8a).

**Figure 3.**
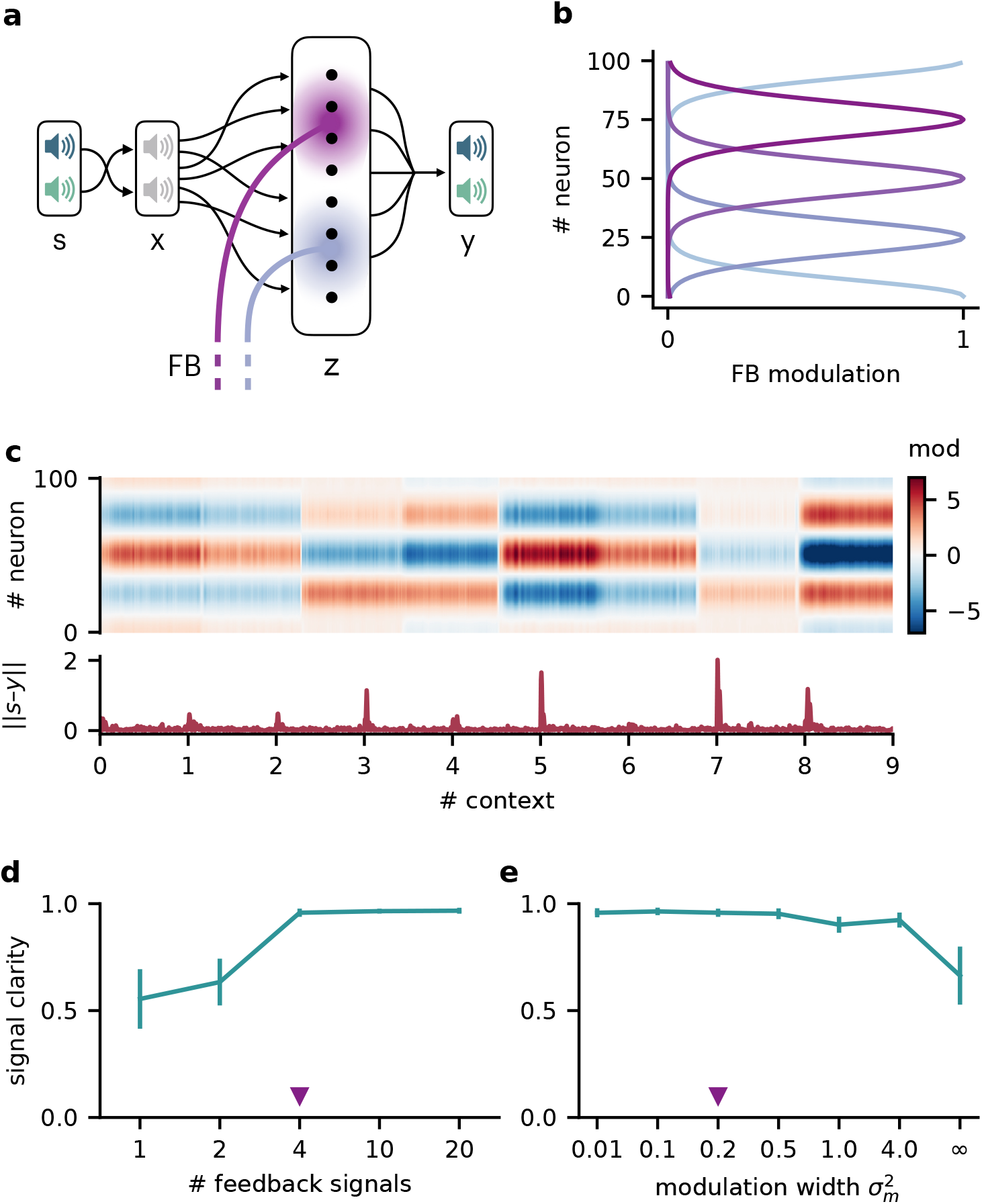
Feedback modulation in the model can be spatially diffuse. **a.** Schematic of the feedforward network with a population that receives diffuse feedback-driven modulation. **b.** Spatial spread of the modulation mediated by 4 modulatory feedback signals with a width of 0.2. **c.** Top: Per neuron modulation during 8 different contexts. Bottom: Corresponding deviation of the network output from sources. **d.** Mean signal clarity across 20 contexts for different numbers of feedback signals; modulation width is 0.2. Error bars indicate standard deviation. Purple triangle indicates default parameters used in (c). **e.** Same as (d) but for different modulation widths; number of feedback signals is 4. The modulation width “∞” corresponds to uniform modulation across the population.

This population-based model with less specific feedback modulation could still solve the dynamic blind source separation task. The diffuse feedback modulation switched when the context changed, but was roughly constant within contexts (Fig. 3c), as in the simple model. The effective weight from the stimuli to the network output also inverted the linear mixture of the sources (Supp. Fig. S8d, cf. Fig. 1c).

We found that only a few distinct feedback signals were needed for a clean separation of the sources across contexts (Fig. 3d). Moreover, the feedback could have a spatially broad effect on the modulated population without degrading the signal clarity (Fig. 3e, Supp. Fig. S8), consistent with the low dimensionality of the context.

We conclude that, in our model, neuromodulation does not need to be spatially precise to enable flexible processing. Given that the suggested feedback-driven modulation mechanism works for slow and diffuse feedback signals, it could in principle be realised by neuromodulatory pathways present in the brain.

### Invariance emerges at the population level

Having established that slow and spatially diffuse feedback modulation enables context-invariant processing, we next investigated the underlying mechanisms at the single neuron and population level. Given that the readout of the population activity was fixed, it is not clear how the context-dependent modulation of single neurons could give rise to a context-independent network output. One possible explanation is that some of the neurons neurons are context-invariant and are exploited by the readout. However, a first inspection of neural activity indicated that single neurons are strongly modulated by context (Fig. 4a). To quantify this, we determined the signal clarity for each neuron at each stage of the feedforward network, averaged across contexts (Fig. 4b). As expected, the signal clarity was low for the sensory stimuli. Intriguingly, the same was true for all neurons of the modulated neural population, indicating no clean separation of the sources at the level of single neurons. Although most neurons had a high signal clarity in some of the contexts, there was no group of neurons that consistently represented one or the other source (Fig. 4c). Furthermore, the average signal clarity of the neurons did not correlate with their contribution to the readout (Fig. 4d). Since single neuron responses were not invariant, context invariance must arise at the population level.

**Figure 4.**
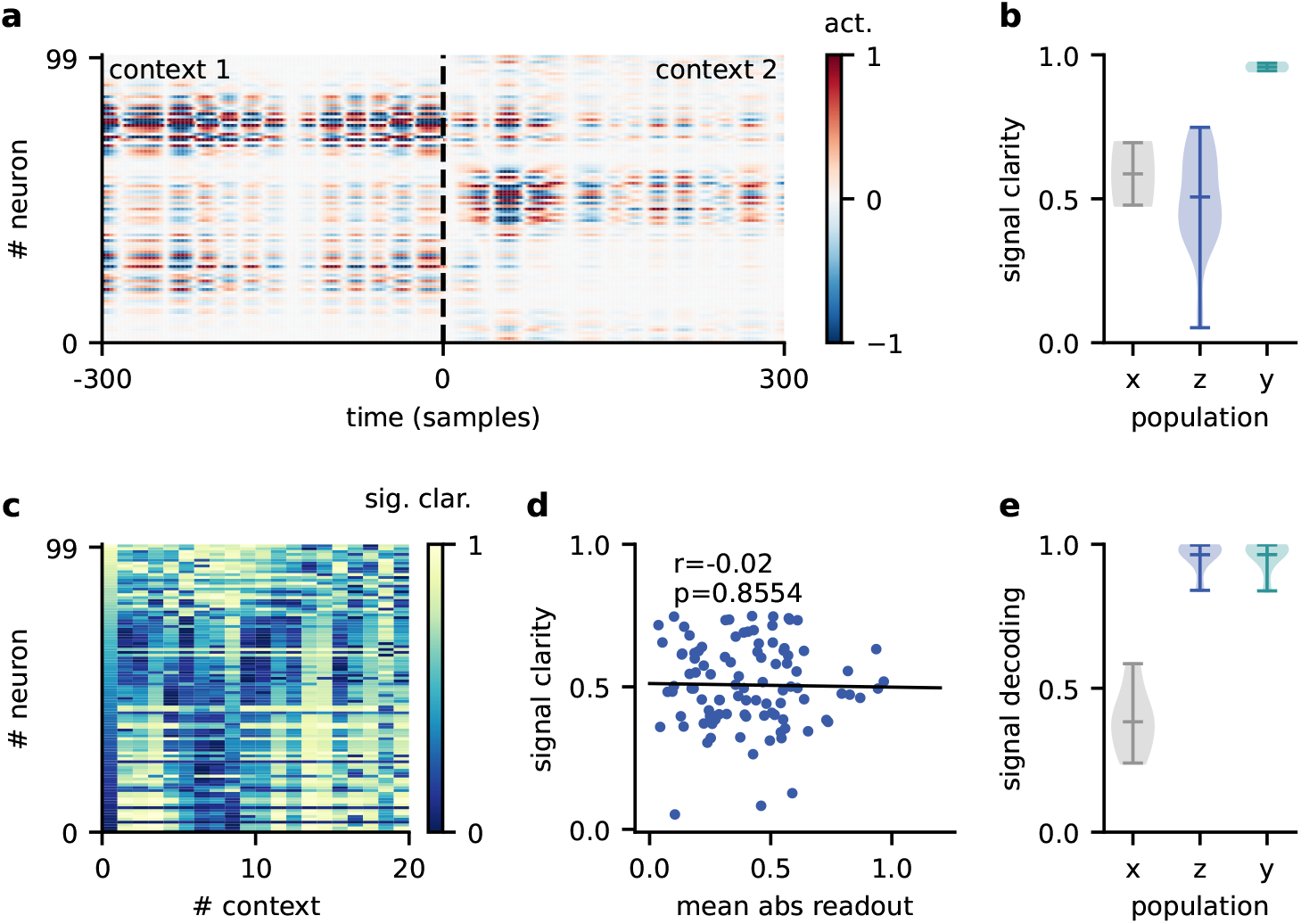
Invariance emerges at the population level. **a.** Population activity in two contexts. **b.** Violin plot of the signal clarity in the sensory stimuli (*x*), neural population (*z*), and network output (*y*), computed across 20 different contexts. **c.** Signal clarity of single neurons in the modulated population for different contexts. **d.** Correlation between average signal clarity over contexts and magnitude of neurons’ readout weight. Corresponding Pearson *r* and *p*-value are indicated in the panel. **e.** Violin plot of the linear decoding performance of the sources from different stages of the feedforward network, computed across 20 contexts. The decoder was trained on a different set of 20 contexts.

To confirm this, we asked how well the sources could be decoded at different stages of the feedforward network. We trained a single linear decoder of the sources on one set of contexts and tested its generalisation to novel contexts. We found that the decoding performance was poor for the sensory stimuli (Fig. 4e), indicating that these did not contain a context-invariant representation. In contrast, the sources could be decoded with high accuracy from the modulated population.

This demonstrates that while individual neurons were not invariant, the population activity contained a context-invariant subspace. In fact, the population had to contain an invariant subspace, because the fixed linear readout of the population was able to extract the sources across contexts. However, the linear decoding approach shows that this subspace can be revealed from the population activity itself with only a few contexts and no knowledge of how the neural representation is used downstream. The same approach could therefore be used to reveal context-invariant subspaces in neural data from population recordings. Note, that the learned readout and the decoder obtained from population activity are not necessarily identical, due to the high dimensionality of the population activity compared to the sources.

### Feedback re-orients the population representation

The question remains how exactly the context-invariant subspace is maintained by feedback modulation. In contrast to a pure feedforward model of invariant perception (Kriegeskorte,2015; Yamins and DiCarlo, 2016), feedback-mediated invariance requires time to establish after contextual changes. Experimentally, hallmarks of this adaptive process should be visible when comparing the population representations immediately after a change and at a later point in time. Our model allows to cleanly separate the early and the late representation by freezing the feedback signals in the initial period after a contextual change (Fig. 5a), thereby disentangling the effects of feedback and context on population activity.

**Figure 5.**
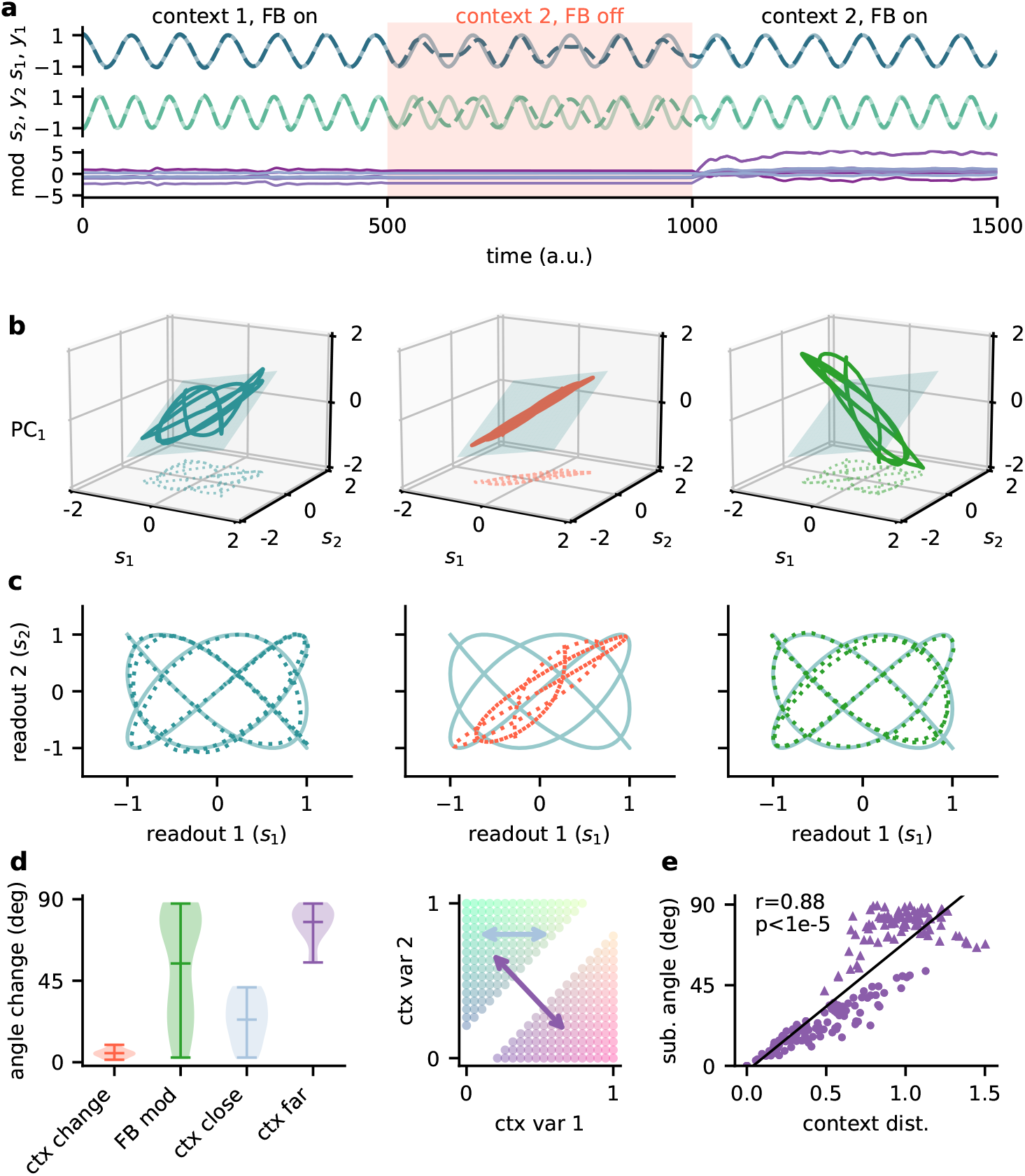
Feedback re-orients the population representation. **a.** Network output (top) and feedback modulation (bottom) for two contexts. The feedback modulation is frozen for the initial period after the context changes. **b.** Population activity in the space of the two readout axes and the first principal component. Projection onto the readout is indicated in the bottom (see (c)). The signal representation is shown for different phases of the experiment. Left: context 1 with intact feedback, center: context 2 with frozen feedback, right: context 2 with intact feedback. The blue plane spans the population activity subspace in context 1 (left). **c.** Same as (b), but projected onto the readout space (dotted lines in (b)). The light blue trace corresponds to the sources. **d.** Left: Change in subspace orientation across 40 repetitions of the experiment, measured by the angle between the original subspace and the subspace for context changes (ctx change), feedback modulation (FB mod) and feedback modulation for similar contexts (ctx close) or dissimilar contexts (ctx far). Right: two-dimensional context space, defined by the coefficients in the mixing matrix. Arrows indicate similar (light blue) and dissimilar contexts (purple). **e.** Distance between pairs of contexts versus the angle between population activity subspaces for these contexts. Circles indicate similar contexts (from the same side of the diagonal, see (d)) and triangles dissimilar contexts (from different sides of the diagonal). Pearson *r* and *p*-value indicated in the panel.

The simulated experiment consisted of three stages: First, the feedback was intact for a particular context and the network outputs closely tracked the sources. Second, the context was changed but the feedback modulation was frozen at the same value as before. As expected, this produced deviations of the output from the sources. Third, for the same context the feedback modulation was turned back on, which reinstated the source signals in the output. In this experiment, we used pure sines as signals for visualisation purposes (Fig. 6a, c). To visualise the population activity in the three stages of the experiment, we considered the space of the two readout dimensions and the first principal component (Fig. 5b). We chose this space rather than, e.g., the first three principal components (Supp. Fig. S9), because it provides an intuitive illustration of the invariant subspace.

**Figure 6.**
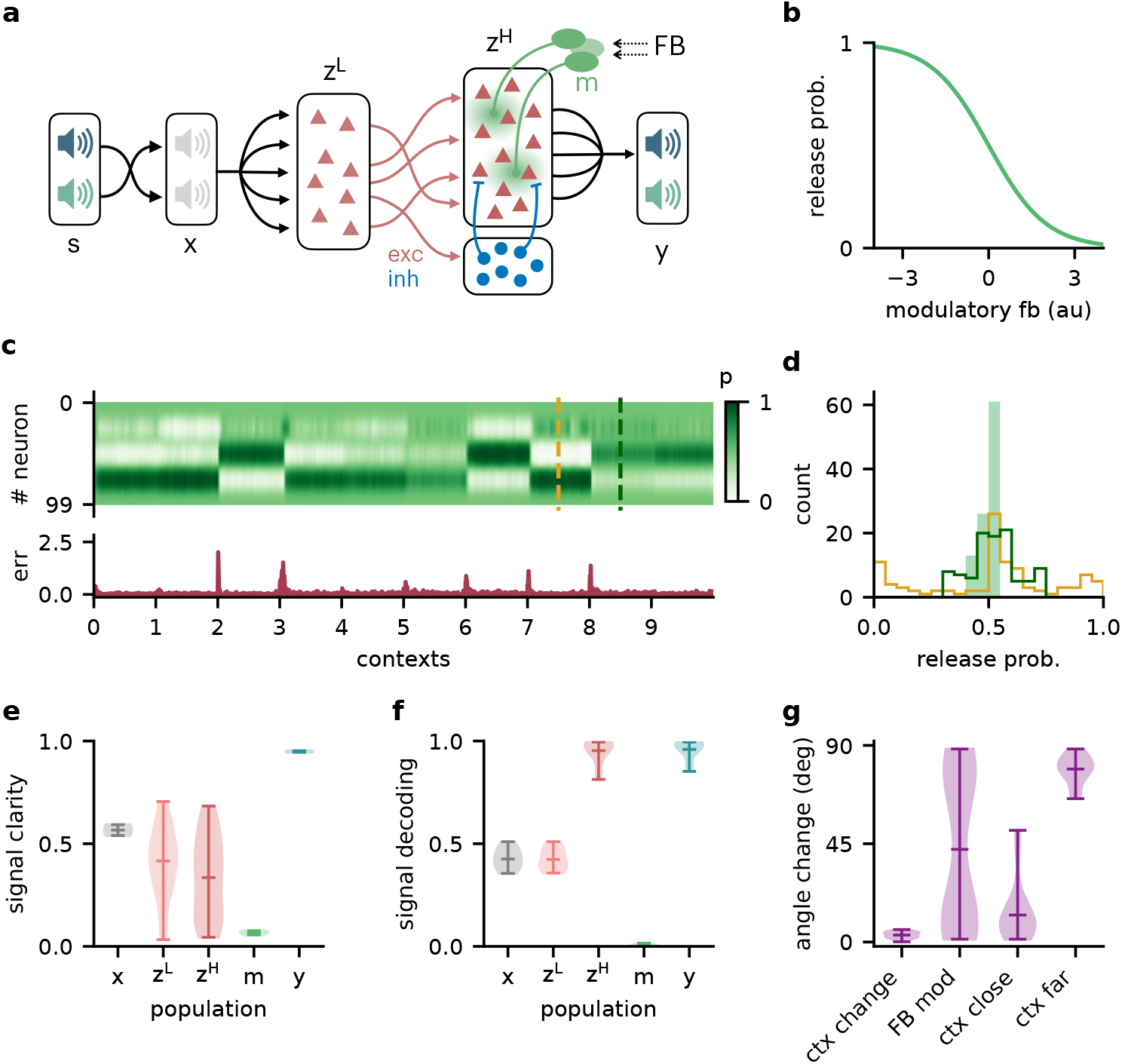
Feedback-driven gain modulation in a hierarchical rate network. **a.** Schematic of the Dalean network comprising a lower- and higher-level population (*z*^L^ and *z*^H^), a population of local inhibitory neurons (blue) and diffuse gain modulation mediated by modulatory interneurons (green). **b.** Decrease in gain (i.e. release probability) with stronger modulatory feedback. **c.** Top: Modulation of neurons in the higher-level population for 10 different contexts. Bottom: Corresponding deviation of outputs *y* from sources *s*. **d.** Histogram of neuron-specific release probabilities averaged across 20 contexts (filled, light green) and during two different contexts (yellow & dark green, see (c)). **e.** Violin plot of signal clarity at different stages of the Dalean model: sensory stimuli (*x*), lower-level (*z*^L^) and higher-level population (*z*^H^), modulatory units (*m*) and network output (*y*), computed across 20 contexts (cf. Fig. 4a). **f.** Violin plot of linear decoding performance of the sources from the same stages as in (e) (cf. Fig. 4d). **g.** Feedback modulation re-orients the population activity (cf. Fig. 5d).

Because the sources were two-dimensional, the population activity followed a pattern within a two-dimensional subspace (Fig. 5b, left; Supp. Fig. S9a). For intact feedback, this population activity matched the sources when projected onto the readout (Fig. 5c, left). Changing the context while freezing the feedback rotated and stretched this representation within the same subspace, such that the readout did not match the sources (Fig. 5b & c, center). Would turning the feedback modulation back on simply reverse this transformation to re-establish an invariant subspace? We found that this was not the case. Instead, the feedback rotated the representation out of the old subspace (Fig. 5b, right), thereby re-orienting it into the invariant readout (Fig. 5c, right).

To quantify the transformation of the population representation, we repeated this experiment multiple times and determined the angle between the neural subspaces. Consistent with the illustration in Fig. 4b, changing the context did not change the subspace orientation, whereas unfreezing the feedback caused a consistent re-orientation (Fig. 4d). The magnitude of this subspace re-orientation depended on the similarity of the old and new context. Similar contexts generally evoked population activity with similar subspace orientations (Fig. 5d,e). This highlights that there is a consistent mapping between contexts and the resulting low-dimensional population activity.

In summary, the role of feedback-driven modulation in our model is to re-orient the population representation in response to changing contexts such that an invariant subspace is preserved.

### The mechanism generalises to a hierarchical Dalean network

So far, we considered a linear network, in which neural activity could be positive and negative. Moreover, feedback modulation could switch the sign of the neurons’ downstream influence, which is inconsistent with Dale’s principle. We wondered if the same population-level mechanisms would operate in a Dalean network, in which feedback is implemented as a positive gain modulation. Although gain modulation is a broadly observed phenomenon that is attributed to a range of cellular mechanisms (Ferguson and Cardin, 2020; Salinas and Thier, 2000), its effect at the population level is less clear (Shine et al., 2021).

We extended the feedforward model as follows (Fig. 6a): First, all neurons had positive firing rates. Second, we split the neural population (*z* in the previous model) into a “lower-level” (*z*^L^) and “higher-level” population (*z*^H^). The lower-level population served as a neural representation of the sensory stimuli, whereas the higher-level population was modulated by feedback. This allowed a direct comparison between a modulated and an unmodulated neural population. It also allowed us to include Dalean weights between the two populations. Direct projections from the lower-level to the higher-level population were excitatory. In addition, a small population of local inhibitory neurons provided feedforward inhibition to the higher-level population. Third, the modulation of the higher-level population was implemented as a local gain modulation that scaled the neural responses. As a specific realisation of gain modulation, we assumed that feedback targeted inhibitory interneurons (e.g., in layer 1; Abs et al., 2018; Ferguson and Cardin, 2020; Malina et al., 2021) that mediate the modulation in the higher-level population (e.g., via presynaptic inhibition; Pardi et al., 2020; Naumann and Sprekeler, 2020). This means that stronger feedback decreased the gain of neurons (Fig. 4b). We will refer to these modulatory interneurons as modulation units *m* (green units in Fig. 4a).

We found that this biologically more constrained model could still learn the context-invariant processing task (Supp. Fig. S10a,b). Notably, the network’s performance did not depend on specifics of the model architecture, such as the target of the modulation or the number of inhibitory neurons (Supp. Fig. S10c-e). In analogy to the previous model, the gain modulation of individual neurons changed with the context and thus enabled the flexible processing required to account for varying context (Fig. 4c). The average gain over contexts was similar across neurons, whereas within a context the gains were broadly distributed (Fig. 4d).

To verify if the task is solved by the same population-level mechanism, we repeated our previous analyses on the single neuron and population level. Indeed, all results generalised to the Dalean network with feedback-driven gain modulation (cf. Fig. 4, 5 & 6). Single neurons in the higher- and lower-level population were not context-invariant (Fig. 6e), but the higher-level population contained a context-invariant subspace (Fig. 6f). This was not the case for the lower-level population, underscoring that invariant representations do not just arise from projecting the sensory stimuli into a higher dimensional space. Instead, the invariant subspace in the higher-level population was again maintained by the feedback modulation, which re-oriented the population activity in response to context changes (Fig. 6g).

### Feedback conveys a non-linear representation of the context

Since single neurons in the higher-level population were not invariant to context, the population representation must also contain contextual information. Indeed, contextual variables could be linearly decoded from the higher-level population activity (Fig. 7a). In contrast, decoding the context from the lower-level population gave much lower accuracy. This shows that the contextual information is not just inherited from the sensory stimuli but conveyed by the feedback via the modulatory units. We therefore expected that the modulatory units themselves would contain a representation of the context. To our surprise, decoding accuracy on the modulatory units was low. This seems counter-intuitive, especially since the modulatory units clearly co-varied with the contextual variables (Fig. 7b). To understand these seemingly conflicting results, we examined how the context was represented in the activity of the modulation units. We found that the modulation unit activity did encode the contextual variables, albeit in a nonlinear way (Fig. 7c). The underlying reason is that the feedback modulation needs to remove contextual variations, which requires nonlinear computations. Specifically, the blind source separation task requires an inversion of the linear mixture of sources. Consistent with this idea, non-linear decoding approaches performed better (Fig. 7d). In fact, the modulatory units contained a linear representation of the “inverse context” (i.e., the inverse mixing matrix, see Methods and Models).

**Figure 7.**
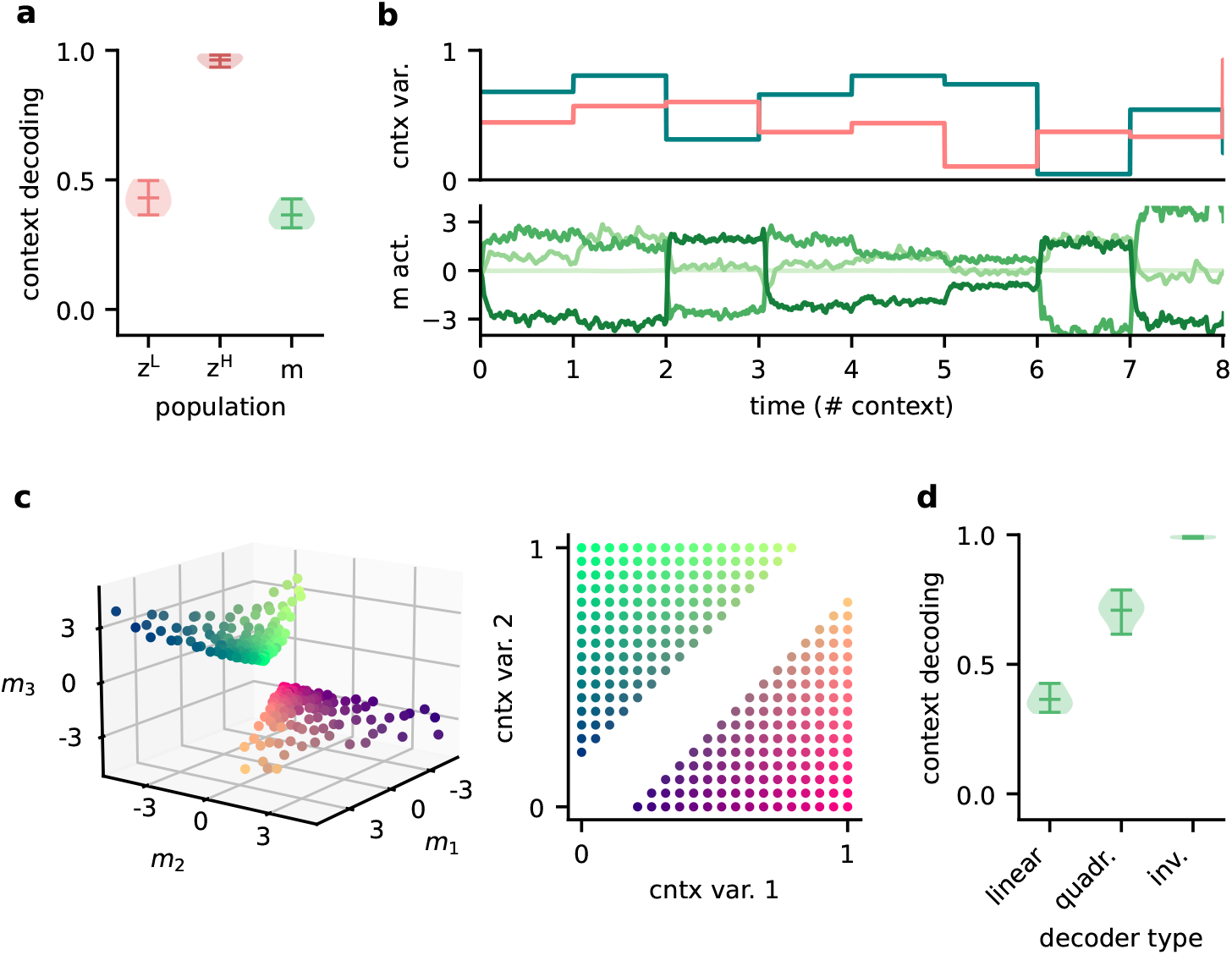
Feedback conveys a non-linear representation of the context. **a.** Linear decoding performance of the context (i.e. mixing) from the network. **b.** Context variables (e.g. source locations, top) and activity of modulatory interneurons (bottom) over contexts; one of the modulatory interneurons is silent in all contexts. **c.** Left: Activity of the three active modulatory interneurons (see b) for different contexts. The context variables are colour-coded as indicated on the right. **d.** Performance of different decoders trained to predict the context from the modulatory interneuron activity. Decoder types are a linear decoder, a decoder on a quadratic expansion and a linear decoder trained to predict the inverse of the mixing matrix.

In summary, the higher-population provides a linear representation not only of the stimuli, but also of the context. In contrast, the modulatory units contained a nonlinear representation of the context, which could not be extracted by linear decoding approaches. We speculate that if contextual feedback modulation is mediated by interneurons in layer 1, they should represent the context in a nonlinear way.

## Discussion

Accumulating evidence suggests that sensory processing is strongly modulated by top-down feedback projections (Gilbert and Li, 2013; Keller and Mrsic-Flogel, 2018). Here, we demonstrate that feedback-driven gain modulation of a feedforward network could underlie stable perception in varying contexts. The feedback can be slow, spatially diffuse and low-dimensional. To elucidate how the context invariance is achieved, we performed single neuron and population analyses. We found that invariance was not evident at the single neuron level, but only emerged in a subspace of the population representation. The feedback modulation dynamically transformed the manifold of neural activity patterns such that this subspace was maintained across contexts. Our results provide further support that gain modulation at the single cell level enables non-trivial computations at the population level (Failor et al., 2021; Shine et al., 2021).

### Invariance in sensory processing

As an example of context-invariant sensory processing, we chose a dynamic variant of the blind source separation task. This task is commonly illustrated by a mixture of voices at a cocktail party (Cherry, 1953; McDermott, 2009). For auditory signals, bottom-up mechanisms of frequency segregation can provide a first processing step for the separation of multiple sound sources (Bronkhorst, 2015; McDermott, 2009). However, separating more complex sounds requires additional active top-down processes (Parthasarathy et al., 2020; Oberfeld and Kloeckner-Nowotny, 2016). In our model top-down feedback guides the source separation itself, while the selection of a source would occur at a later processing stage – consistent with recent evidence for “late selection”(Brodbeck et al., 2020; Yahav and Golumbic, 2021).

Although blind source separation is commonly illustrated with auditory signals, the suggested mechanism of context-invariant perception is not limited to a given sensory modality. The key nature of the task is that it contains stimulus dimensions that need to be encoded (the sources) and dimensions that need to be ignored (the context). In visual object recognition, for example, the identity of visual objects needs to be encoded, while contextual variables such as size, location, orientation, or surround need to be ignored. Neural hallmarks of invariant object recognition are present at the population level (DiCarlo and Cox, 2007; DiCarlo et al., 2012; Hong et al., 2016), and to some extent also on the level of single neurons (Quiroga et al., 2005). Classically, the emergence of invariance has been attributed to the extraction of invariant features in feedforward networks (Riesenhuber and Poggio, 1999; Wiskott and Sejnowski, 2002; DiCarlo and Cox, 2007; Kriegeskorte, 2015), but recent work also highlights the role of recurrence and feedback (Gilbert and Li, 2013; Kar et al., 2019; Kietzmann et al., 2019). Here, we focused on the role of feedback, but clearly, feedforward and feedback processes are not mutually exclusive and likely work in concert to create invariance. Their relative contribution to invariant perception requires further studies and may depend on the invariance in question.

Similarly, how invariance can be learned will depend on the underlying mechanism. The feedback-driven mechanism we propose is reminiscent of meta-learning consisting of an inner and an outer loop (Hochreiter et al., 2001; Wang et al., 2018b). In the inner loop, the modulatory system infers the context to modulate the feedforward network accordingly. This process is unsupervised. In the outer loop, the modulatory system is trained to generalise across contexts. Here, we performed this training using supervised learning, which requires the modulatory system to experience the sources in isolation (or at least obtain an error signal). Such an identification of the individual sources could, e.g., be aided by other sensory modalities (McDermott, 2009). However, the optimisation of the modulatory system does not necessarily require supervised learning. It could also be guided by task demands via reinforcement learning, or by unsupervised priors such as a non-Gaussianity of the outputs.

### Mechanisms of feedback-driven gain modulation

There are different ways in which feedback can affect local processing. Here, we focused on gain modulation (McAdams and Maunsell, 1999; Reynolds and Heeger, 2009; Vinck et al., 2015). Neuronal gains can be modulated by a range of mechanisms (Ferguson and Cardin, 2020; Shine et al., 2021). In our model, the mechanism needs to satisfy a few key requirements: i) the modulation is not uniform across the population, ii) it operates on a timescale similar to that of changes in context, and iii) it is driven by a brain region that has access to the information needed to infer the context.

Classical neuromodulators such as acetylcholine (Disney et al., 2007; Kawai et al., 2007), dopamine(Thurley et al., 2008) or serotonin (Azimi et al., 2020) are signalled through specialised neuromodulatory pathways from subcortical nuclei (van den Brink et al., 2019). These neuromodulators can control the neural gain depending on behavioural states such as arousal, attention or expectation of rewards (Ferguson and Cardin, 2020; Hasselmo and McGaughy, 2004; Bayer and Glimcher, 2005; Polack et al., 2013; Kuchibhotla et al., 2017). Their effect is typically thought to be brain-wide and long-lasting, but recent advances in measurement techniques (Sabatini and Tian, 2020; Lohani et al., 2020) indicate that it could be area- or even layer-specific, and vary on sub-second time scales (Lohani et al., 2020; Bang et al., 2020; Poorthuis et al., 2013; Pinto et al., 2013).

More specific feedback projections arrive in layer 1 of the cortex, where they target the distal dendrites of pyramidal cells and inhibitory interneurons (Douglas and Martin, 2004; Roth et al., 2016; Marques et al., 2018). Dendritic input can change the gain of the neural transfer function on fast timescales (Larkum et al., 2004; Jarvis et al., 2018). The spatial scale of the modulation will depend on the spatial spread of the feedback projections and the dendritic arbourisation. Feedback to layer 1 interneurons provides an alternative mechanism of local gain control. In particular, neuron-derived neurotrophic factor-expressing interneurons (NDNF) in layer 1 receive a variety of top-down feedback projections and produce GABAergic volume transmission (Abs et al., 2018), thereby down-regulating synaptic transmission (Miller, 1998; Laviv et al., 2010). This gain modulation can act on a timescale of hundreds of milliseconds (Branco and Staras, 2009; Urban-Ciecko et al., 2015;Malina et al., 2021; Molyneaux and Hasselmo, 2002), and, although generally considered diffuse, can also be synapse type-specific (Chittajallu et al., 2013).

The question remains where in the brain the feedback signals originate. Our model requires the responsible network to receive feedforward sensory input to infer the context. In addition, feedback inputs from higher-level sensory areas to the modulatory system allow a better control of the modulated network state. Higher-order thalamic nuclei are ideally situated to integrate different sources of sensory inputs and top-down feedback (Sampathkumar et al., 2021) and mediate the resulting modulation by targeting layer 1 of lower-level sensory areas (Purushothaman et al., 2012; Roth et al., 2016; Sherman, 2016). In our task setting, the inference of the context requires the integration of sensory signals over time and therefore recurrent neural processing. For this kind of task, thalamus may not be the site of contextual inference, because it lacks the required recurrent connectivity (Halassa and Sherman, 2019). However, contextual inference may be performed by higher-order cortical areas, and could either be relayed back via the thalamus or transmitted directly, for example, via cortico-cortical feedback connections.

### Testable predictions

Our model makes several predictions that could be tested in animals performing invariant sensory perception. Firstly, our model indicates that invariance across contexts may only be evident at the neural population level, but not on the single cell level. Probing context invariance at different hierarchical stages of sensory processing may therefore require population recordings and corresponding statistical analyses such as neural decoding (Glaser et al., 2020). Secondly, we assumed that this context invariance is mediated by feedback modulation. The extent to which context invariance is enabled by feedback on a particular level of the sensory hierarchy could be studied by manipulating feedback connections. Since layer 1 receives a broad range of feedback inputs from different sources, this may require targeted manipulations. If no effect of feedback on context invariance is found, this may either indicate that feedforward mechanisms dominate or that the invariance in question is inherited from an earlier stage, in which it may well be the result of feedback modulation. Given that feedback is more pronounced in higher cortical areas (McAdams and Maunsell, 1999; Pardi et al., 2020), we expect that the contribution of feedback may play a larger role for the more complex forms of invariance further up in the sensory processing hierarchy. Thirdly, for feedback to mediate context invariance, the feedback projections need to contain a representation of the contextual variables. Our findings suggest, however, that the detection of this representation may require a non-linear decoding method. Finally, a distinguishing feature of feedback and feedforward mechanisms is that feedback mechanisms take more time. We found that immediately following a sudden contextual change, the neuronal representation initially changes within the manifold associated with the previous context. Later, the feedback reorients the manifold to reestablish the invariance on the population level. Whether these dynamics are a signature of feedback processing or also present in feedforward networks will be an interesting question for future work.

### Comparison to prior work

Computational models have implicated neuronal gain modulation for a variety of functions (Salinas and Sejnowski, 2001; Reynolds and Heeger, 2009). Even homogeneous changes in neuronal gain can achieve interesting population effects (Shine et al., 2021), such as orthogonalisation of sensory responses (Failor et al., 2021). More heterogeneous gain modulation provides additional degrees of freedom that enables, for example, attentional modulation (Reynolds and Heeger, 2009; Carandini and Heeger, 2012), coordinate transformations (Salinas and Thier, 2000) and – when amplified by recurrent dynamics – a rich repertoire of neural trajectories (Stroud et al., 2018). Gain modulation has also been suggested as a means to establish invariant processing (Salinas and Abbott, 1997), as a biological implementation of dynamic routing (Olshausen et al., 1993). While the modulation in these models of invariance can be interpreted as an abstract form of feedback, the resulting effects on the population level were not studied.

An interesting question is by which mechanisms the appropriate gain modulation is computed. In previous work, gain factors were often learned individually for each context, for example by gradient descent or Hebbian plasticity (Olshausen et al., 1993; Salinas and Abbott, 1997; Stroud et al., 2018), mechanisms that may be too slow to achieve invariance on a perceptual timescale (Wiskott, 2006). In our model, by contrast, the modulation is dynamically controlled by a recurrent network. Once it has been trained, such a recurrent modulatory system can rapidly infer the current context, and provide an appropriate feedback signal on a timescale only limited by the modulatory mechanism.

### Limitations and future work

In our model, we simplified many aspects of sensory processing. Using simplistic sensory stimuli – compositions of sines – allowed us to focus on the mechanisms at the population level, while avoiding the complexities of natural sensory stimuli and deep sensory hierarchies. Although we do not expect conceptual problems in generalising our results to more complex stimuli, such as speech or visual stimuli, the associated computational challenges are substantial. For example, the feedback in our model was provided by a recurrent network, whose parameters were trained by back-propagating errors through the network and through time. This training process can get very challenging for large networks and long temporal dependencies (Bengio et al., 1994; Pascanu et al., 2013).

In our simulations we trained the whole model – the modulatory system, the sensory representation and the readout. For the simplistic stimuli we used, we observed that the training process mostly concentrated on optimising the modulatory system and readout, while a random mapping of sensory stimuli to neural representations seemed largely sufficient to solve the task. For more demanding stimuli, we expect that the sensory representation the modulatory system acts upon may become more important. A well-suited representation could minimise the need for modulatory interventions (Finn et al., 2017), in a coordinated interaction of feedforward and feedback.

To understand the effects of feedback modulation on population representations, we included biological constraints in the feedforward network and the structure of the modulatory feedback. However, we did not strive to provide a biologically plausible implementation for the computation of the appropriate feedback signals, and instead used an off-the-shelf recurrent neural network (Hochreiter and Schmidhuber, 1997). The question how these signals could be computed in a biologically plausible way remains for future studies. The same applies to the question how the appropriate feedback signals can be learned by local learning rules (Lillicrap et al., 2020) and how neural representations and modulatory systems learn to act in concert.

## Methods and Models

To study how feedback-driven modulation can enable flexible sensory processing, we built models of feedforward networks that are modulated by feedback. The feedback was dynamically generated by a modulatory system, which we implemented as a recurrent network. The weights of the recurrent network were trained such that the feedback modulation allowed the feedforward network to solve a flexible invariant processing task.

### The dynamic blind source separation task

As an instance of flexible sensory processing we used a dynamic variant of blind source separation. In classical blind source separation, two or more unknown time-varying sources 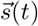 need to be recovered from a set of observations (i.e. sensory stimuli) 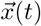. The sensory stimuli are composed of an unknown linear mixture of the sources such that 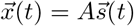 with a fixed mixing matrix *A*. Recovering the sources requires to find weights *W* such that 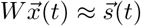. Ideally, *W* is equal to the pseudo-inverse of the unknown mixing matrix *A*, up to permutations.

In our dynamic blind source separation task, we model variations in the stimulus context by changing the linear mixture over time – albeit on a slower timescale than the time-varying signals. Thus, the sensory stimuli are constructed as

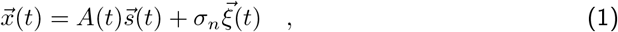

where *A*(*t*) is a time-dependent mixing matrix and *σ_n_* is the amplitude of additive white noise 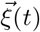. The time-dependent mixing matrix determines the current context and was varied in discrete time intervals *n_t_*, meaning that the mixing matrix *A*(*t*) (i.e. the context) was constant for *n_t_* samples before it changed. The goal of the dynamic blind source separation task is to recover the original signal sources 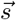 from the sensory stimuli 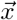 across varying contexts. Thus, the network model output needs to be invariant to the specific context of the sources. Note that while the context was varied, the sources themselves were the same throughout the task, unless stated otherwise. Furthermore, in the majority of experiments the number of source signals and sensory stimuli was *n_s_* = 2. A list of default parameters for the dynamic blind source separation task can be found in Table 1.

**Table 1.** Default parameters of the dynamic blind source separation task

| parameter | symbol | value |
| --- | --- | --- |
| number of signals | $n_s$ | 2 |
| number of samples in context | $n_t$ | 1000 |
| additive noise | $\sigma_n$ | 0.001 |
| sampling frequency | $f_s$ | 8 kHz |

### Source signals

As default source signals we used two compositions of two sines each (“chords”) with a sampling rate of *f_s_* = 8000Hz that can be written as

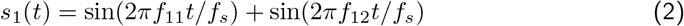

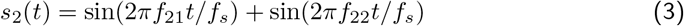

with frequencies *f*_11_ = 100 Hz, *f*_12_ = 125 Hz, *f*_21_ = 150 Hz and *f*_22_ = 210 Hz. Note that in our model we measure time as the number of samples from the source signals, meaning that timescales are relative and could be arbitrarily rescaled.

In Fig 5, we used pure sine signals with frequency *f* for visualisation purposes: *s_i_* = sin(2*πft*/*f_s_*). We also validated the model on signals that are not made of sine waves, as a sawtooth and a square wave signal (Supp. Fig. S4). Unless stated otherwise, the same signals were used for training and testing the model.

### Time-varying contexts

We generated the mixing matrix *A* for each context by drawing random weights from a uniform distribution between 0 and 1, allowing only positive mixtures of the sources. Unless specified otherwise, we sampled new contexts for each training batch and for the test data, such that the training and test data followed the same distribution without necessarily being the same. The dimension of the mixing matrices was determined by number of signals *n_s_* such that A was of shape *n_s_* × *n_s_*. To keep the overall amplitude of the sensory stimuli in a similar range across different mixtures, we normalised the row sums of each mixing matrix to one. In the case of *n_s_* = 2, this implies that the contexts (i.e. the mixing matrices) are drawn from a 2-dimensional manifold (see Fig. 8, bottom left). In addition, we only used the randomly generated mixing matrices whose determinant was larger than some threshold value. We did this to ensure that each signal mixture was invertible and that the weights needed to invert the mixing matrix were not too extreme. A threshold value of 0.2 was chosen based on visual inspection of the weights from the inverted mixing matrix.

**Figure 8.**
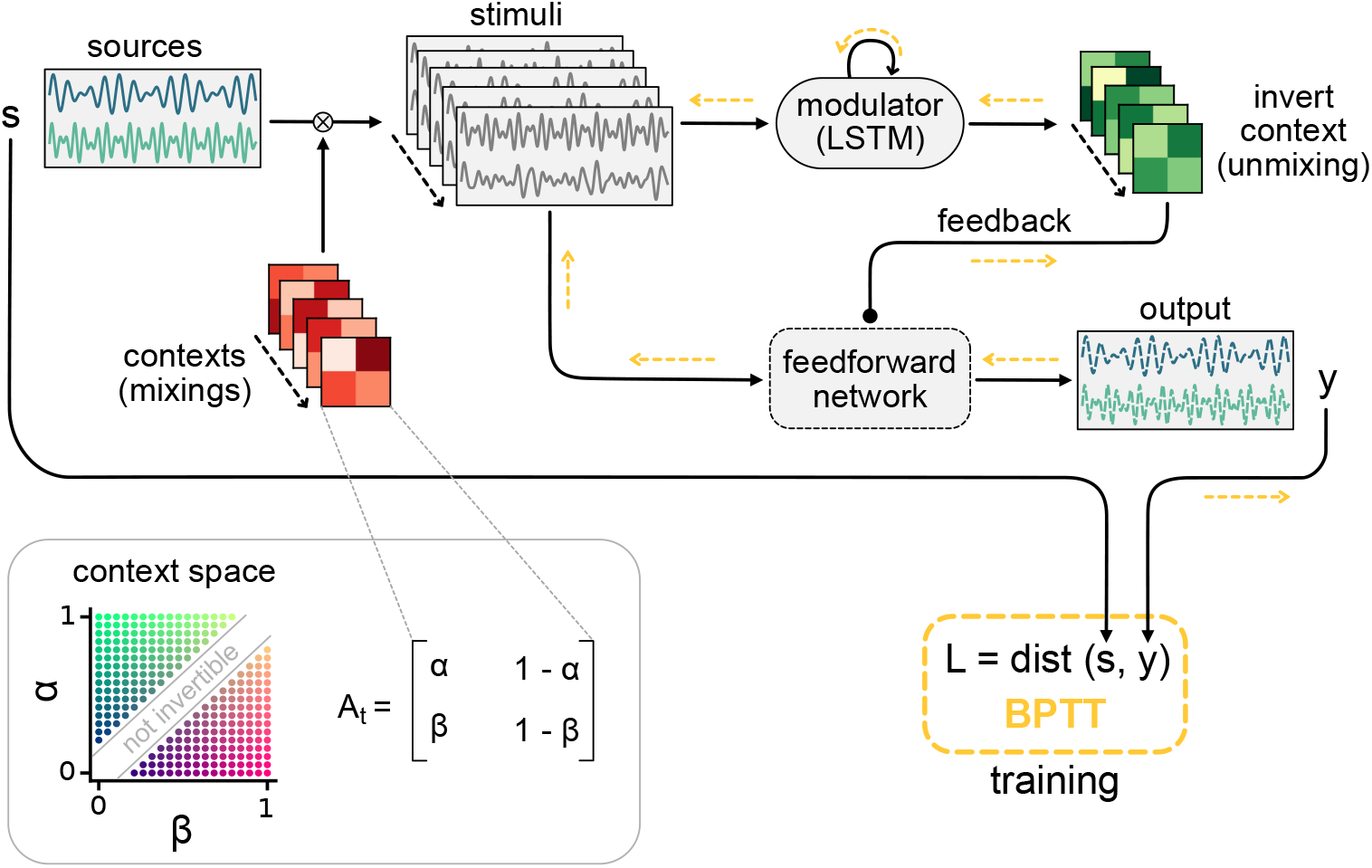
Schematic of the dynamic blind source separation task, the context space and the modulated feedforward network. Information flow is indicated by black arrows and the flow of the error during training with backpropagation through time (BPTT) is shown in yellow.

### Modulated feedforward network models

Throughout this work, we modelled feedforward networks of increasing complexity. Common to all networks was that they received the sensory stimuli 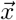 and should provide an output 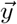 that matches the source signals 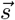. In the following, we first introduce the simplest model variant and how it is affected by feedback from the modulatory system, and subsequently describe the different model extensions.

### Modulation of feedforward weights by a recurrent network

In the simplest feedforward network the network output 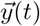 is simply a linear readout the sensory stimuli 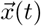, with readout weights that are dynamically changed by the modulatory system:

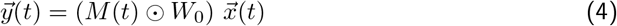

where *W*_0_ are the baseline weights and *M*(*t*) the modulation provided by the modulatory system. *M*(*t*) is of the same shape as *W*_0_ and determines the element-wise multiplicative modulation of the baseline weights. Because the task requires the modulatory system to dynamically infer the context, we modelled it as a recurrent network – more specifically a long-short term memory network (LSTMs; Hochreiter and Schmidhuber, 1997) – with *N_h_* = 100 hidden units. In particular, we used LSTMs with forget gates (Gers et al., 2000) but no peephole connections (for an overview of LSTM variants see Greff et al. (2016)).

In this work we treated the LSTM as a black-box modulatory system that receives the sensory stimuli and the feedforward network’s output and provides the feedback signal in return (Fig. 1a). A linear readout of the LSTM’s output determines the modulation *M*(*t*) in Eq. (4). In brief, this means that

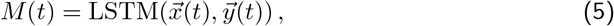

where LSTM(·) is a function that returns the LSTM readout. For two-dimensional sources and sensory stimuli, for instance, LSTM(·) receives a concatenation of the two-dimensional vectors 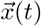 and 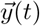 as input and returns a two-by-two feedback modulation matrix – one multiplicative factor for each weight in *W*_0_. The baseline weights *W*_0_ were randomly drawn from the Gaussian distribution 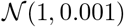 and fixed throughout the task. The LSTM parameters and readout were learned during training of the model.

### Extension 1: Reducing the temporal specificity of feedback modulation

To probe our model’s sensitivity to the timescale of the modulatory feedback (Fig. 2), we added a temporal filter to Eq. (5). In that case the modulation *M*(*t*) followed the dynamics

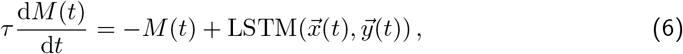

with *τ* being the time constant of modulation. For small *τ*, the feedback rapidly affects the feedforward network, whereas larger *τ* imply a slowly changing modulatory feedback signal. The unit of this timescale is the number of samples from the source signals. Note that the timescale of the modulation should be considered relative to the timescale of the context changes *n_t_*. As a default time constant we used *τ* = 100 < *n_t_* (see Table 2).

**Table 2.** Default parameters of the network models

| parameter | symbol | value |
| --- | --- | --- |
| number of hidden units in LSTM | $N_h$ | 100 |
| number of units in middle layer $z$ | $N_z$ | 100 |
| number of distinct feedback signals | $N_{FB}$ | 4 |
| number of neurons in lower-level population | $N_L$ | 40 |
| number of neurons in higher-level population | $N_H$ | 100 |
| number of inhibitory neurons | $N_I$ | 20 |
| timescale of modulation | $\tau$ | 100 |
| spatial spread of modulation | $\sigma_m^2$ | 0.2 |

### Extension 2: Reducing the spatial specificity of feedback modulation

To allow for spatially diffuse feedback modulation (Fig. 3), we added an intermediate layer between the sensory stimuli and the network output. This intermediate layer consisted of a population of *N_z_* = 100 units that were modulated by the feedback, where neighbouring units were modulated similarly. More specifically, the units were arranged on a ring to allow for a spatially constrained modulation without boundary effects. The population’s activity vector 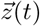 is described by

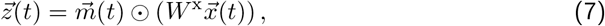

with the sensory stimuli 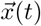, a weight matrix *W^x^* of size *N_z_* × *n_s_* and the vector of unit-specific multiplicative modulations 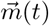. Note that the activity of the units was not constrained to be positive here. The output of the network was then determined by a linear readout of the population activity vector according to

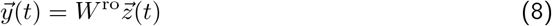

with a fixed readout matrix *W*^ro^.

The modulation to a single unit *i* was given by

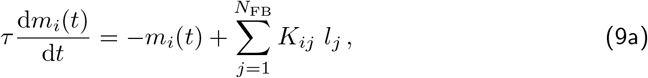

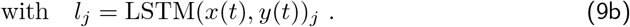

Here, *τ* is the modulation time constant, *K* a kernel that determines the spatial specificity of modulation, LSTM(·)_*j*_ the *j*-th feedback signal from the LSTM and *N*_FB_ the total number of feedback signals. As in the simple model, the *N*_FB_ feedback signals were determined by a linear readout from LSTM.

The modulation kernel *K* was defined as a set of von Mises functions:

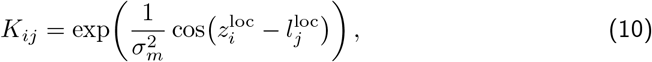

where 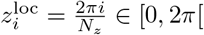 represents the location of the modulated unit *i* on the ring and 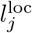 the ‘‘preferred location” of modulatory unit *j*, i.e., the location on the ring that it modulates most effectively. These ‘preferred locations” 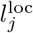 of the feedback units were evenly distributed on the ring. The variance parameter 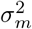 determines the spatial spread of the modulatory effect of the feedback units, i.e., the spatial specificity of the modulation. Overall, the spatial distribution of the modulation was therefore determined by the number of distinct feedback signals *N*_FB_ and their spatial spread 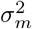 (see Table 2 for a list of network parameters).

### Extension 3: Hierarchical rate-based network

We further extended the model with spatial modulation (Eqs. (7)–(10)) to include a two-stage hierarchy, positive rates and synaptic weights that obey Dale’s law. Furthermore, we implemented the feedback modulation as a gain modulation that scales neural rates but keeps them positive. To this end, we modelled the feedforward network as a hierarchy of a lower-level and a higher-level population. Only the higher-level population received feedback modulation. Splitting the neural populations in this way allowed us to model the connections between them with weights that follow Dale’s law. Furthermore, the unmodulated lower-level population could serve as a control for the emergence of context-invariant representations. The lower-level population consisted of *N*_L_ = 40 rate-based neurons and the population activity vector was given by

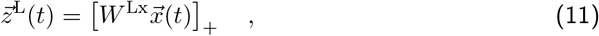

where *W*^Lx^ is a fixed weight matrix, 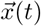 the sensory stimuli and the rectification [·]_+_ = max(0, ·) ensures that rates are positive. The lower-level population thus provides a neural representation of the sensory stimuli. The higher-level population consisted of *N*_H_ = 100 rate-based neurons that received feedforward input from the lower-level population. The feedforward input consisted of direct excitatory projections as well as feedforward inhibition through a population of *N*_I_ = 20 local inhibitory neurons. The activity vector of the higher-level population 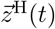 was thus given by

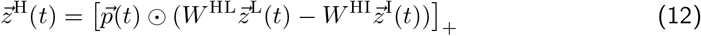

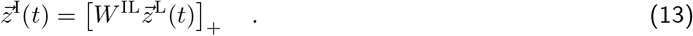

Here *W*^HL^, *W*^HI^ and *W*^IL^ are positive weight matrices, 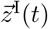 the inhibitory neuron activities and 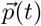 the neuron-specific gain modulation factors. As for the spatially modulated network of Extension 2, the network output 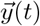 was determined by a fixed linear readout *W*^ro^ (see Eq. (8)). The distributions used to randomly initialise the weight matrices are provided in Table 3.

**Table 3.** Distributions used for randomly initialised weight parameters

| weights | distribution |
| --- | --- |
| $W_0$ | $\mathcal{N}(1, 0.001)$ |
| $W^x$ | $\mathcal{N}(0, 0.5)$ |
| $W^{Lx}$ | $\mathcal{N}(0, 0.5)$ |
| $W^{ro}$ | $\mathcal{N}(0, 0.5)$ |
| $W^{HL}$ | $\mathcal{N}(1, 0.5) \cdot 20/N_H$ |
| $W^{IL}$ | $\mathcal{N}(1, 0.5)/N_I$ |
| $W^{HI}$ | $\mathcal{N}(1, 1) \cdot 20/N_H$ |
| LSTM parameters | $\mathcal{U}(-\sqrt{1/N_H}, \sqrt{1/N_H})$ |
| LSTM readout | $\mathcal{U}(-\sqrt{1/N_{FB}}, \sqrt{1/N_{FB}})$ |

Again, the modulation was driven by feedback from the LSTM, but in this model variant we assumed inhibitory feedback, i.e., stronger feedback signals monotonically decreased the gain. More specifically, we assumed that the feedback signal targets a population of modulation units 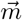, which in turn modulate the gain in the higher-level population. The gain modulation of neuron *i* was constrained between 0 and 1 and determined by

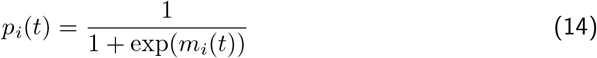

with *m_i_*(*t*) being the activity of a modulation unit *i*, which follows the same dynamics as in Eq. (9a) (see Fig. 6a).

### Training the model

We used gradient descent to find the model parameters that minimise the difference between the source signal 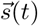 and the feedforward network’s output 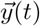:

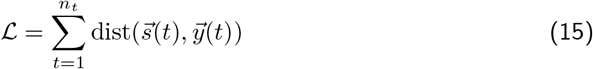

with a distance measure dist(·). We used the machine learning framework PyTorch (Paszke et al., 2019) to simulate the network model, obtain the gradients of the objective 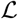 by automatic differentiation and update the parameters of the LSTM using the Adam optimiser (Kingma and Ba, 2014) with a learning rate of *η* = 10^-3^. As distance measure in the objective we used a smooth variant of the L1 norm (PyTorch’s smooth L1 loss variant), because it is less sensitive to outliers than the mean squared error (Huber, 1964).

During training, we simulated the network dynamics over batches of 32 trials using forward Euler with a timestep of Δ*t* = 1. Each trial consisted of *n_t_* time steps (i.e. samples) and the context (i.e. mixing matrix) differed between trials. Since the model contains feedback and recurrent connections, we trained it using backpropagation through time (Werbos, 1990). This means that for each trial, we simulated the model and computed the loss for every time step. At the end of the trial we propagated the error through the *n_t_* steps of the model to obtain the gradients and updated the parameters accordingly (Fig. 8). Although the source signals were the same in every trial, we varied their phase independently across trials to prevent the LSTM from learning the exact signal sequence. To this end, we generated 16,000 samples of the source signals and in every batch randomly selected chunks of *n_t_* samples independently from each source. Model parameters were initialised according to the distributions listed in Table 3.

In all model variants we optimised the parameters of the modulator (input, recurrent and readout weights as well as the biases of the LSTM; see Eq. (5) & (9b)). The parameters were initialised with the defaults from the corresponding PyTorch modules, as listed in Table 3. To facilitate the training in the hierarchical rate-based network despite additional constraints, we also optimised the feedforward weights *W*^HL^, *W*^HI^, *W*^IL^, *W*^Lx^ and *W*^ro^. In principle, this allows to adapt the representation in the two intermediate layers such that the modulation is most effective. However, although we did not quantify it, we observed that optimising the network readout *W*^ro^ facilitated the training the most, suggesting that a specific format of the sensory representations was not required for an effective modulation.

To prevent the gain modulation factor from saturating at 0 or 1, we added a regularisation term 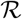 to the loss function Eq. (15) that keeps the LSTM’s output small:

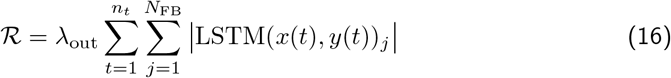

with λ_out_ = 10^-5^.

Gradient values were clipped between −1 and 1 before each update to avoid large updates. For weights that were constrained to be positive, we used their absolute value in the model. Each network was trained for 10,000 to 12,000 batches and for 5 random initialisations (Supp. Fig. S2).

### Testing and manipulating the model

We tested the network model performance on an independent random set of contexts (i.e. mixing matrices), but with the same source signals as during training. During testing, we also changed the context every *n_t_* steps, but the length of this interval was not crucial for performance (Supp. Fig. S2d).

To manipulate the feedback modulation in the hierarchical rate-based network (see Fig. 4), we provided an additional input to the modulation units *m* in Eq. (9a). We used an input of 3 or – 3 depending on whether the modulation units were activated or inactivated, respectively. To freeze the feedback modulation (see Fig. 6), we discarded the feedback signal and held the local modulation *p* in Eq. (14) at a constant value determined by the feedback before the manipulation.The dynamics of the LSTM were continued, but remained hidden to the feedforward network until the freezing was stopped.

### Unmodulated feedforward network models

#### Linear regression

As a control, we trained feedforward networks with weights that were not changed by a modulatory system. First, we used the simplest possible network architecture, in which the sensory stimuli are linearly mapped to the outputs (Fig. S1a):

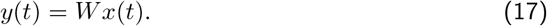

It is intuitive that a fixed set of weights *W* cannot invert two different contexts (i.e. different mixing matrices *A*_1_ and *A*_2_). As an illustration we trained this simple feedforward network on one context and tested it on different contexts. To find the weights *W*, we used linear regression to minimise the mean squared error between the source signal *s*(*t*) and the network’s output *y*(*t*). The training data consisted of 1024 consecutive time steps of the sensory stimuli for a fixed context, and the test data consisted of different 1024 time steps generated under a potentially different mixing. We repeated this procedure by training and testing a network for all combinations of 20 random contexts.

#### Multi-layer nonlinear network

Since solving the task was not possible with a single set of readout weights, we extended the feedforward model to include 3 hidden layers consisting of 32, 16 and 8 rectified linear units (Fig. S1d). The input to this network was one time point from the sensory stimuli and the target output the corresponding time point of the sources. We trained the multi-layer network on 5000 batches of 32 contexts using Adam (learning rate 0.001) to minimise the mean squared error between the network output and the sources.

#### Multi-layer network with sequences as input

Solving the task requires the network to map the same sensory stimulus to different outputs depending on the context. However, inferring the context takes more than one time point. To test if a feedforward network with access to multiple time points at once could in principle solve the task, we changed the architecture of the multi-layer network, such that it receives a sequence of the sensory stimuli (Fig. S1g). The output of the network was a sequence of equal length. We again trained this network on 5000 batches of 32 contexts to minimise the error between its output and the target sources, where both the network input and output were sequences. The length of these sequences was varied between 1 and 150.

### Data analysis

#### Signal clarity

To determine task performance, we measured how clear the representation of the source signals is in the network output. We first computed the correlation coefficient of each signal *s_i_* with each output *y_j_*

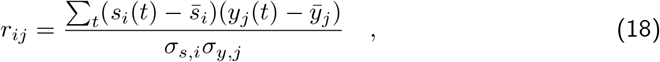

where 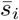 and *ȳ_j_* are the respective temporal mean and *σ_s,i_* and *σ_y,j_* the respective temporal standard deviations. The signal clarity in output *y_j_* is then given by the absolute difference between the absolute correlation with one compared to the other signal:

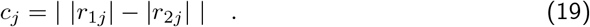

By averaging over outputs we determined the overall signal clarity within the output. Note that the same measure can be computed on other processing stages of the feedforward network. For instance, we used the signal clarity of sources in the sensory stimuli as a baseline control.

### Signal-to-noise ratio

The signal-to-noise ratio in the sensory stimuli was determined as the variability in the signal compared to the noise. Since the mean of both the stimuli and the noise were zero, the signal-to-noise ratio could be computed by

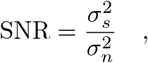

where *σ_n_* was the standard deviation of the additive white noise and *σ_s_* the measured standard deviation in the noise-free sensory stimuli, which was around 0.32. As a scale of the signal-to-noise ratio we used decibels (dB), i.e., we used dB = 10log_10_(SNR).

### Linear decoding analysis

#### Signal decoding

We investigated the population-level invariance by using a linear decoding approach. If there was an invariant population subspace, the source signals could be decoded by the same decoder across different contexts. We therefore performed linear regression between the activity in a particular population and the source signals. This linear decoder was trained on *n_c_* = 10 different contexts with *n_t_* = 1, 000 time points each, such that the total number of samples was 10, 000. The linear decoding was then tested on 10 new contexts and the performance determined using the R^2^ measure.

#### Context decoding

We took a similar approach to determine from which populations the context could be decoded. For the dynamic blind source separation task the context is given by the source mixture, as determined by the mixing matrix. Since we normalised the rows of each mixing matrix, the context was determined by two context variables. We calculated the temporal average of the neuronal activities within each context and performed a linear regression of the context variables onto these averages. To exclude onset transients, we only considered the second half (500 samples) of every context. Contexts were sampled from the two-dimensional grid of potential contexts. More specifically, we sampled 20 points along each dimension and excluded contexts, in which the sensory stimuli were too similar (analogously to the generation of mixing matrices), leaving 272 different contexts (see Fig. 5g, right). The linear decoding performance was determined with a 5-fold cross-validation and measured using R-squared. Since the modulatory feedback signals depend non-linearly on the context (Fig. 5g), we tested two non-linear versions of the decoding approach. First, we performed a quadratic expansion of the averaged population activity before a linear decoding. Second, we tested a linear decoding of the inverse mixing matrix (four weights) instead of the two variables determining the context.

### Population subspace analysis

We visualised the invariant population subspaces by projecting the activity vector onto the two readout dimensions and the first principal component. To measure how the orientation of the subspaces changes when the context or feedback changes, we computed the angle between the planes spanned by the respective subspaces. These planes were fitted on the three-dimensional data described above using the least squares method. Since we were only interested in the relative orientation of the subspaces, we used a circular measure of the angles, such that a rotation of 180 degrees corresponded to 0 degrees. This means that angles could range between 0 and 90 degrees.

## Code availability

The code for models and data analysis will be made available under https://github.com/sprekelerlab/feedback_modulation_Naumann21 upon publication of the project.

## Acknowledgments

We thank Owen Mackwood for providing a code framework that manages simulations on a compute cluster, Loreen Hertäg and Johannes Letzkus for feedback on the manuscript, and the members of the Sprekeler lab for valuable discussions.

## Supplemental Information

**Figure S1.**
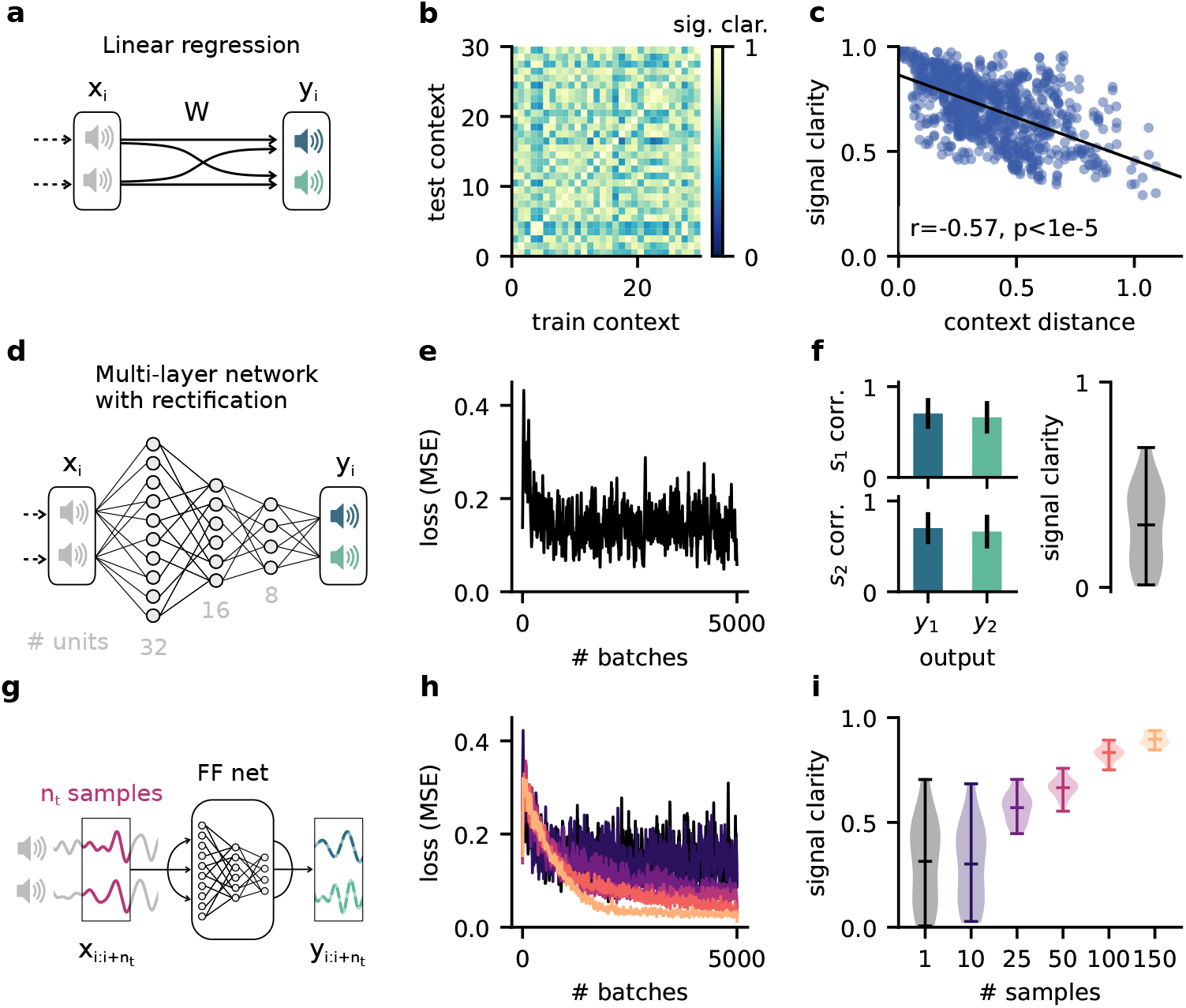
The dynamic blind source separation task cannot be solved with a feedforward network, unless the network receives a sequence of inputs at once. This would require an additional mechanism to retain information over time. **a.** Schematic of a feedforward network consisting of a linear readout only. **b.** Pairwise signal clarity of one context when the network is trained on another context. **c.** Correlation between the distance between two contexts and their pairwise signal clarity (see (b)). **d.** Schematic of a multi-layer feedforward network with three hidden layers (32, 16 and 8 rectified linear units). **e.** Loss during training for the network in (d), measure by the mean squared error between the output and the sources. **f.** Network performance after training. Left: Correlation of the outputs with the sources over 20 contexts. Error bars indicate standard deviation. Right: Signal clarity across 20 contexts for the trained network. **g.** Schematic of network architecture and training setup when using a sequence of *n_t_* samples as input to the multi-layer network. **h.** Same as (e) but for different number of samples. Color code corresponds to (i). **i.** Signal clarity for trained networks that receive different numbers of samples as input.

**Figure S2.**
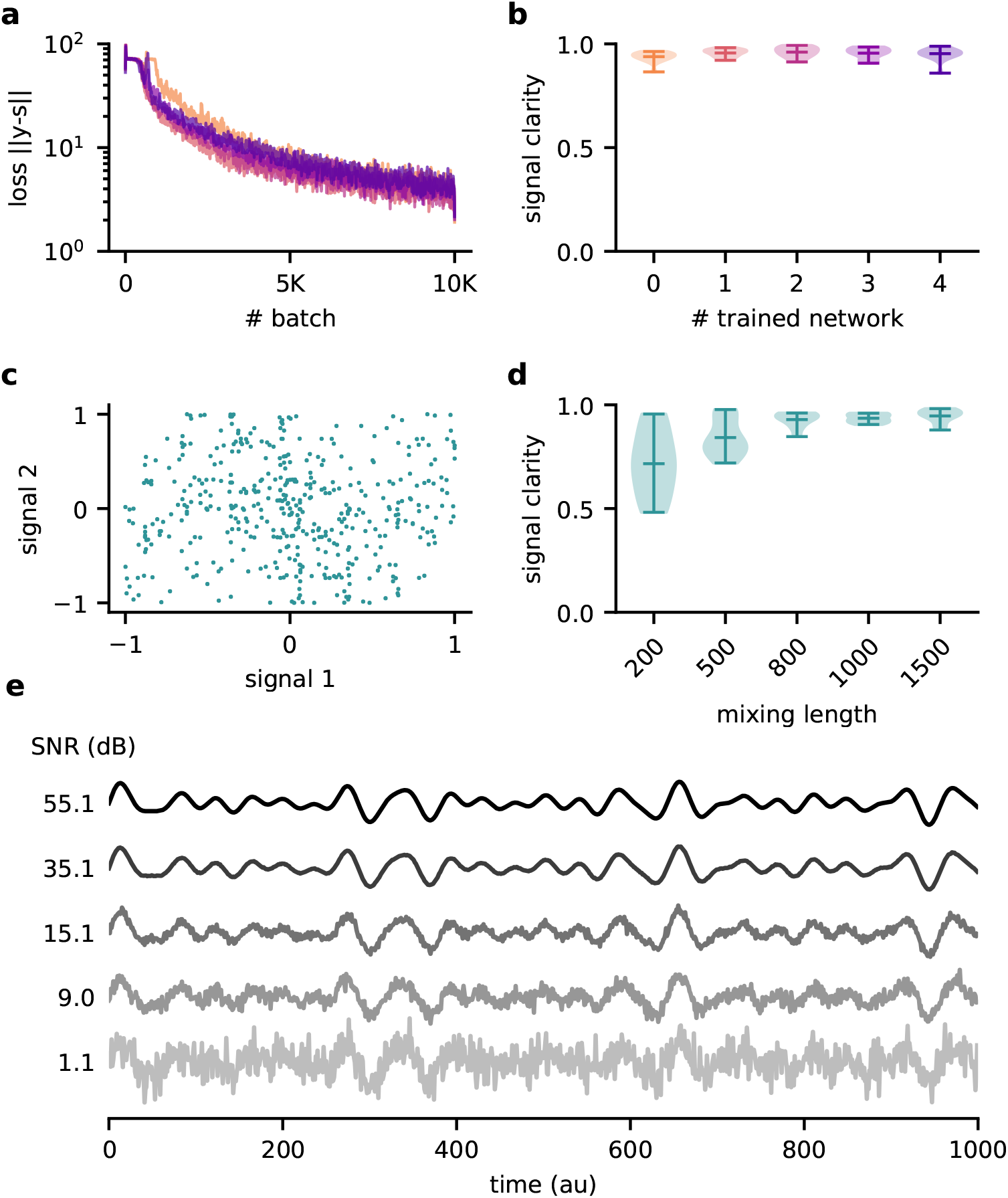
Robustness of the feedback-driven modulation mechanism. **a.** Loss over training for 5 different random initialisations of the model and **b.** signal clarity for 20 test contexts in the corresponding trained networks. The model performance is robust across model instantiations. **c.** Samples from the two default signals are uncorrelated. **d.** Signal clarity for different lengths of the context during testing. The length of the context interval is not crucial for performance, indicating that the network did not learn the interval by heart. **e.** Example traces of the sensory stimuli for different signal-to-noise ratios.

**Figure S3.**
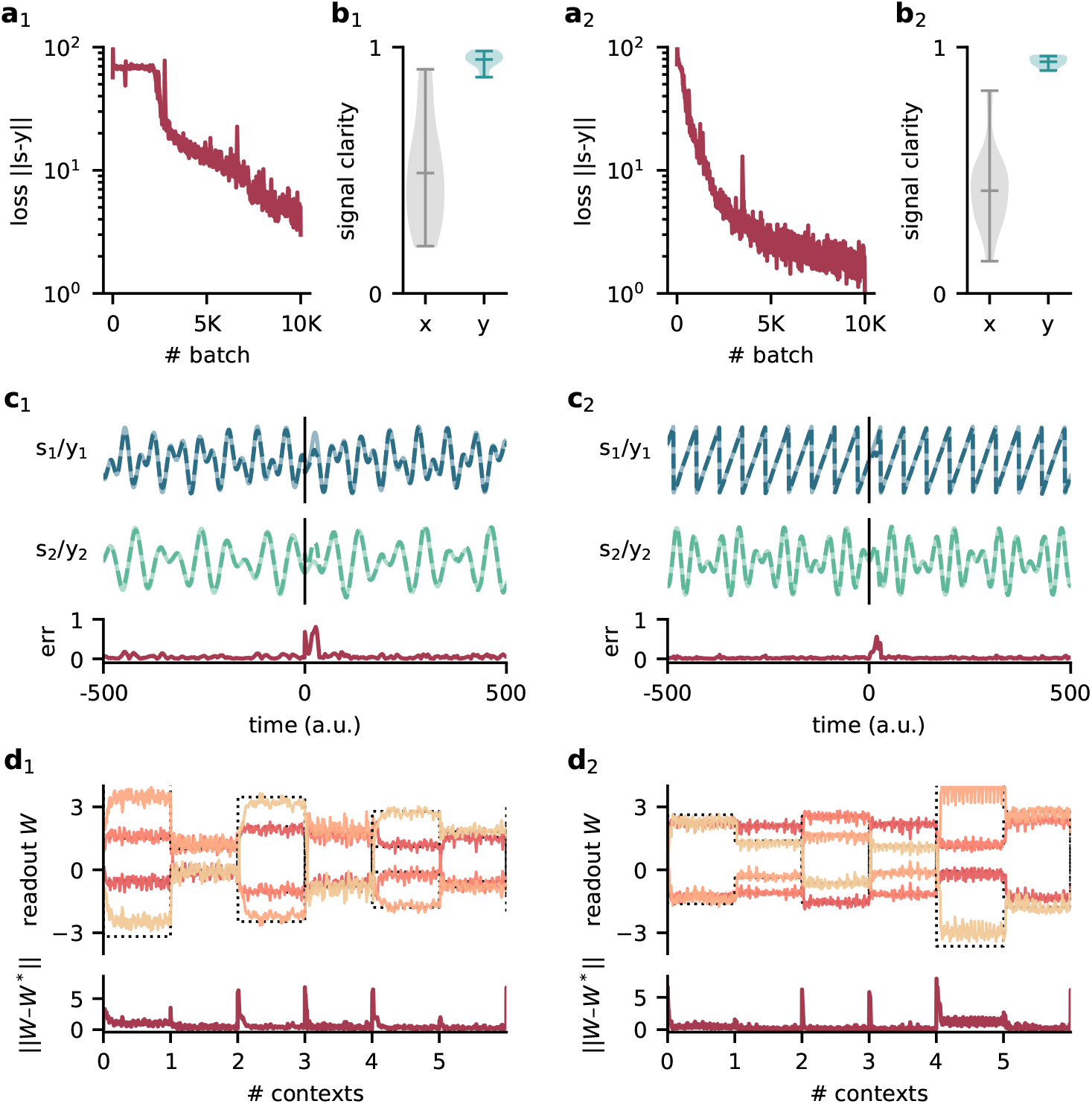
Model performance for two different sets of source signals. Left: Compositions of sines with *f*_11_ = 120 Hz, *f*_12_ = 2.2 Hz, *f*_21_ = 100 Hz and *f*_22_ = 145 Hz. Right: Sawtooth function with frequency 140 Hz and composed sine of 150 Hz and 210 Hz. **a_1/2_**. Loss over training. **b_1/2_**. Signal clarity for 20 test contexts measured in the sensory stimuli and the network output. **c_1/2_**. Example traces of the sources and the network output (top) and corresponding deviation between them (bottom). The context changes at time 0. **d_1/2_**. Top: Readout weights across 6 contexts; dotted lines indicate the optimal weights. Bottom: Deviation of readout from the optimal weights.

**Figure S4.**
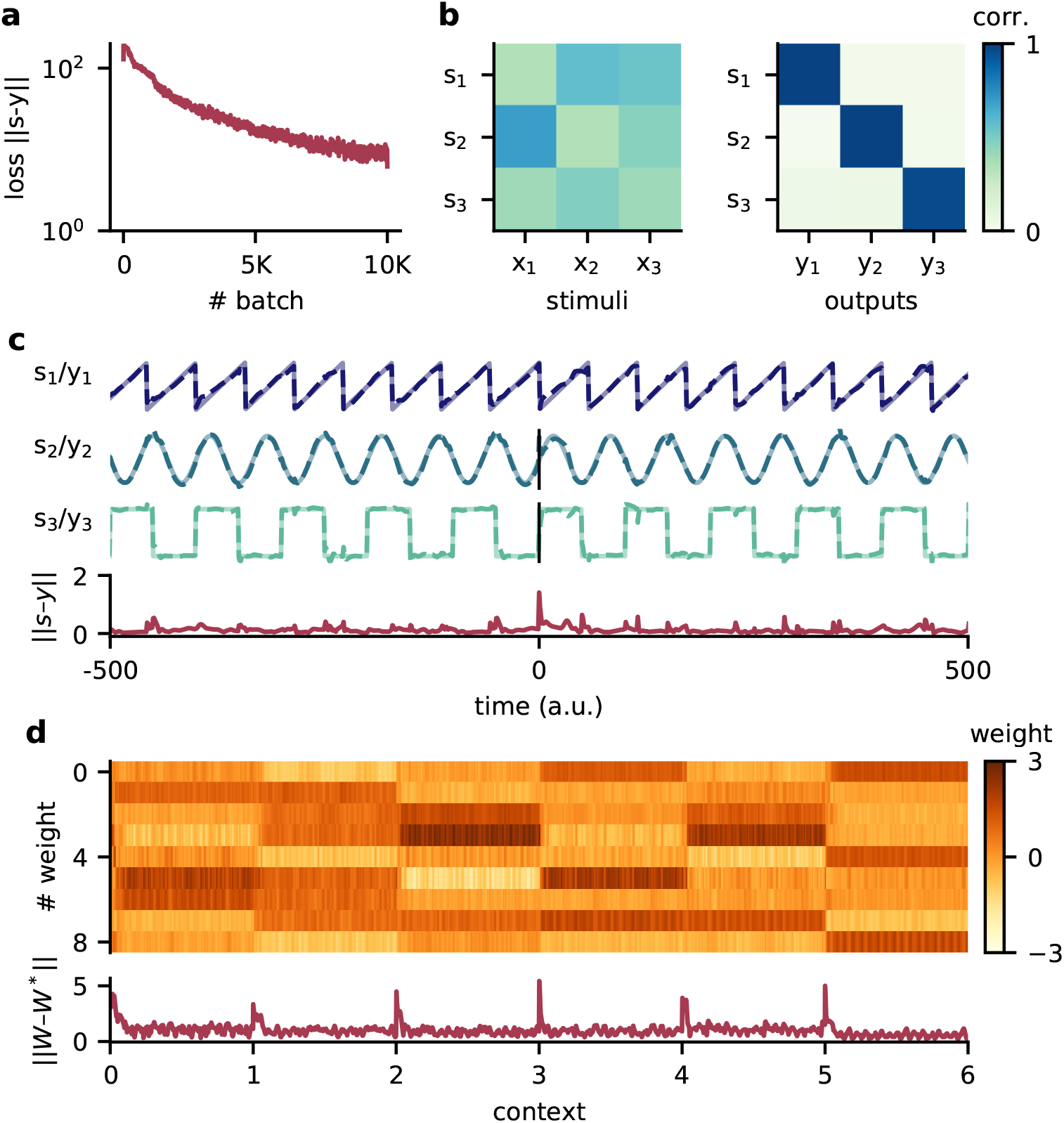
Model performance for three source signals. **a.** Loss over training. **b.** Correlation of the sources with the mixed sensory stimuli (left) and with the network outputs (right). **c.** Example traces of the three source signals and network outputs (top) and corresponding deviation between them (bottom). The context changes at time 0. The source signals are a sawtooth of frequency 140 Hz, a sine wave of frequency 120 Hz and a square wave signal of 80 Hz. **d.** Top: Readout weights across 6 contexts. Bottom: Deviation of readout from the optimal weights.

**Figure S5.**
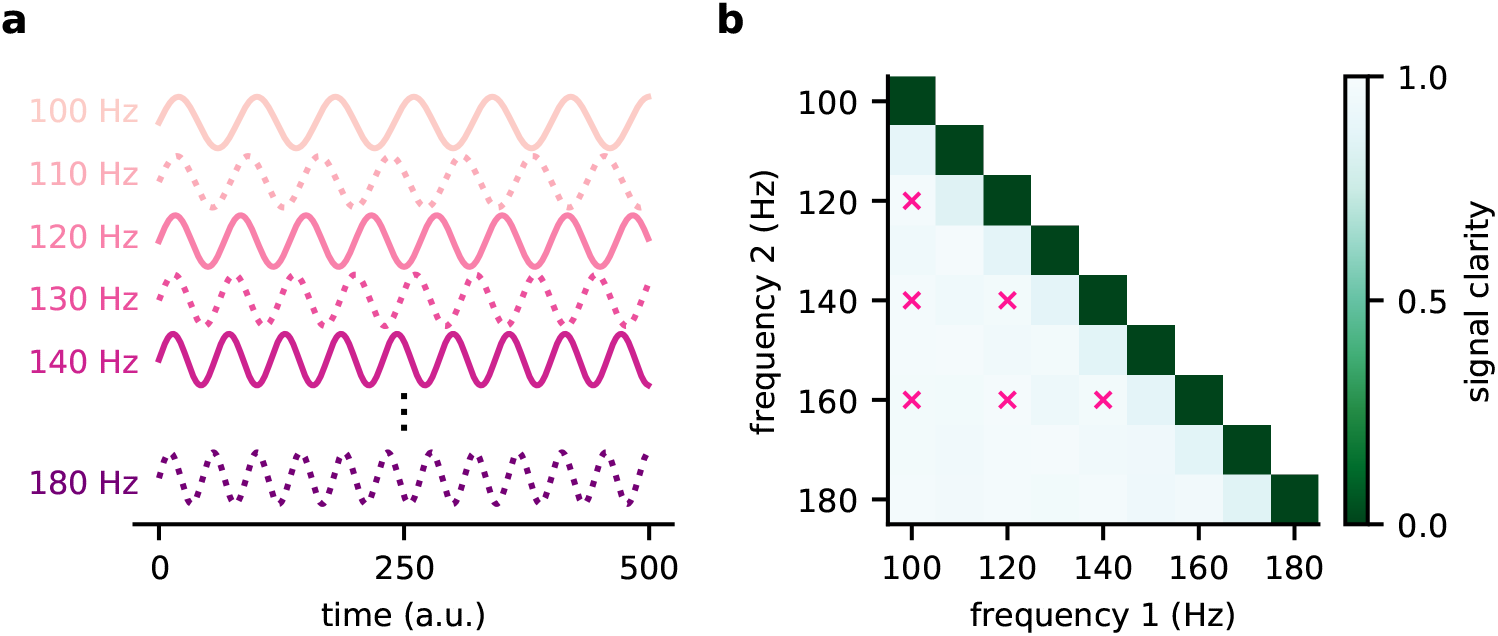
The modulated network model generalises across frequencies. **a.** Illustration of the source signals used during training (solid lines) and only during testing (dotted lines). During the training, the model experiences only a subset of potential signals. **b.** Signal clarity for different combinations of test frequencies. Combinations used during training are marked with a pink cross.

**Figure S6.**
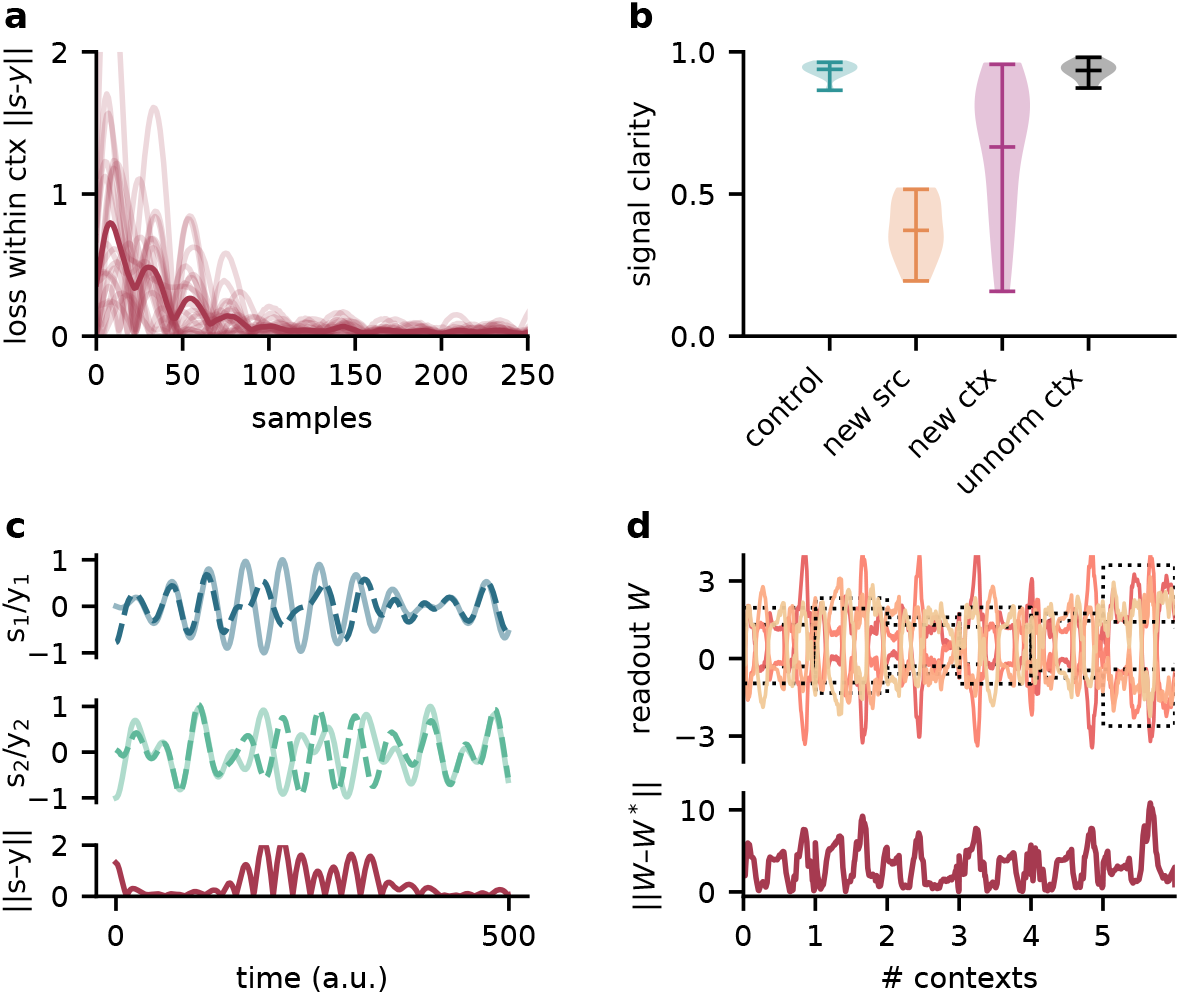
The modulator learns a model of the sources and contexts and infers the current context from the stimuli. Testing the network on sources and contexts with different statistics than during training thus impairs its performance. **a.** Deviation of network output from sources within contexts. Average across contexts shown in dark red. **b.** Signal clarity for different test cases: same sources and same context statistics as during training (‘‘control”), new sources (“new src”), same sources but different context statistic (i.e. unnormed mixing matrices, “new ctx”), and different context statistics but when training the network on them (“unnorm ctx”). **c.** Top: Sources (*s*_1,2_) and network output (*y*_1,2_) for a context when testing on new sources. Bottom: Deviation of outputs from the sources. **d.** Top: Modulated readout weights across 6 contexts when testing on new sources; dotted lines indicate the inverse of the current mixing. Bottom: Deviation of readout from target weights.

**Figure S7.**
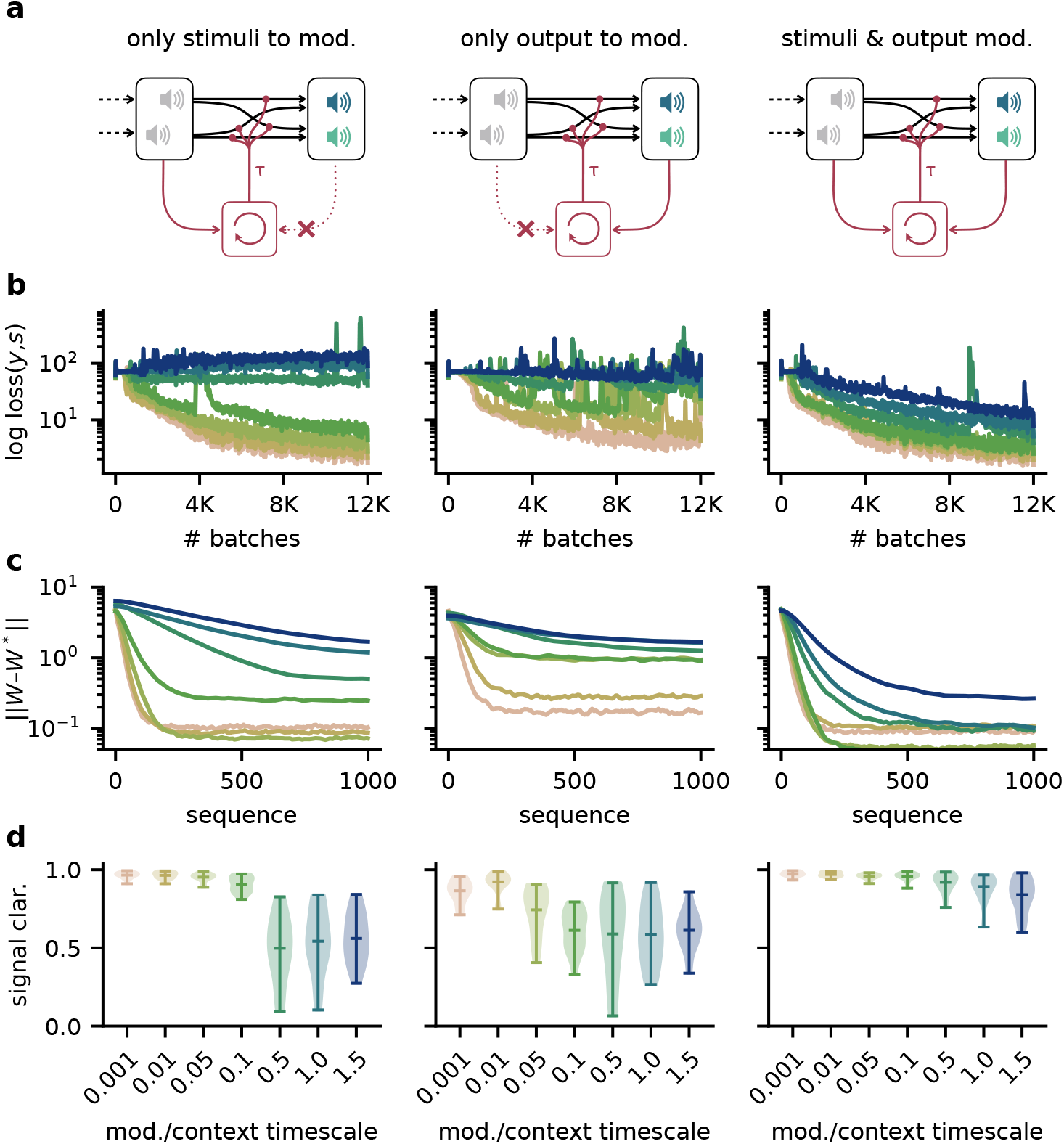
Robustness to slow feedback modulation depends on the inputs to the modulatory system. **a.** Illustration of different input configurations: the modulatory system receives only the sensory stimuli as feedforward input (left), only the network output as feedback input (right) or both (right). **b.** Loss over training for different timescales. Colours correspond to values shown in (d). **c.** Deviation of the readout weights from the optimal weights over the duration of a context for different modulation timescales, averaged across 20 contexts. Colours correspond to values shown in (d). **d.** Signal clarity for different timescales of the modulatory feedback signal.

**Figure S8.**
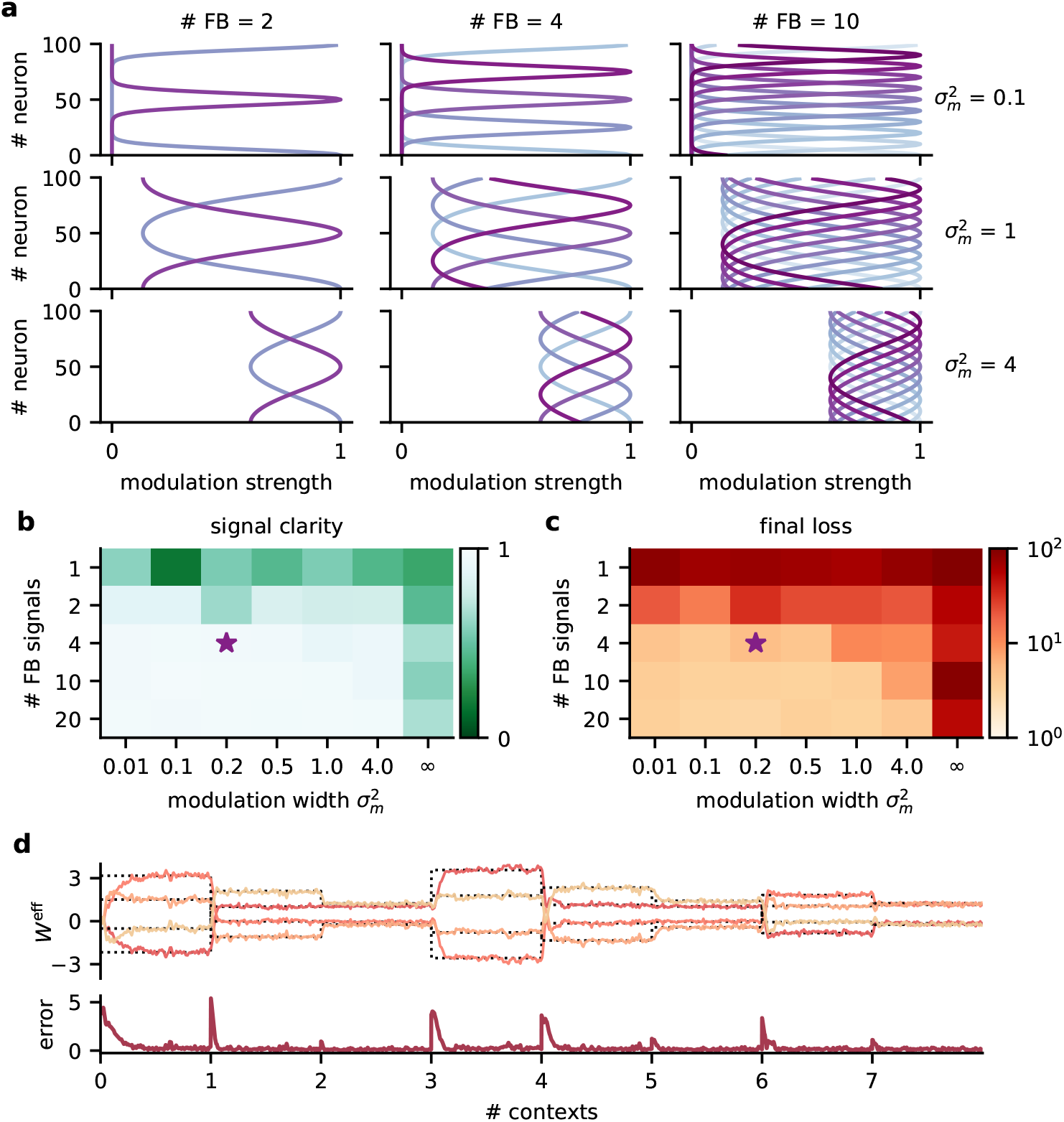
Robustness to the spatial scale of feedback modulation. **a.** Examples of the spatial extent of feedback modulation for different numbers of feedback signals (# FB) and spatial spread 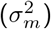. **b.** Signal clarity and **c.** final log loss in network models with different parameters determining the spatial scale of feedback modulation. Signal clarity was averaged across 20 contexts. Final loss was averaged across the last 200 batches during training. The purple star indicates default values used in the main results. Modulation width of “∞” corresponds to a homogeneous modulation over the whole population. **d.** Top: Effective weights from stimuli to network output over 8 contexts. Effective weights are computed as the modulated weights from stimuli to neural population, multiplied with the readout weights. Dotted lines indicate inverse of mixing. Bottom: Deviation of effective weights from the inverse.

**Figure S9.**
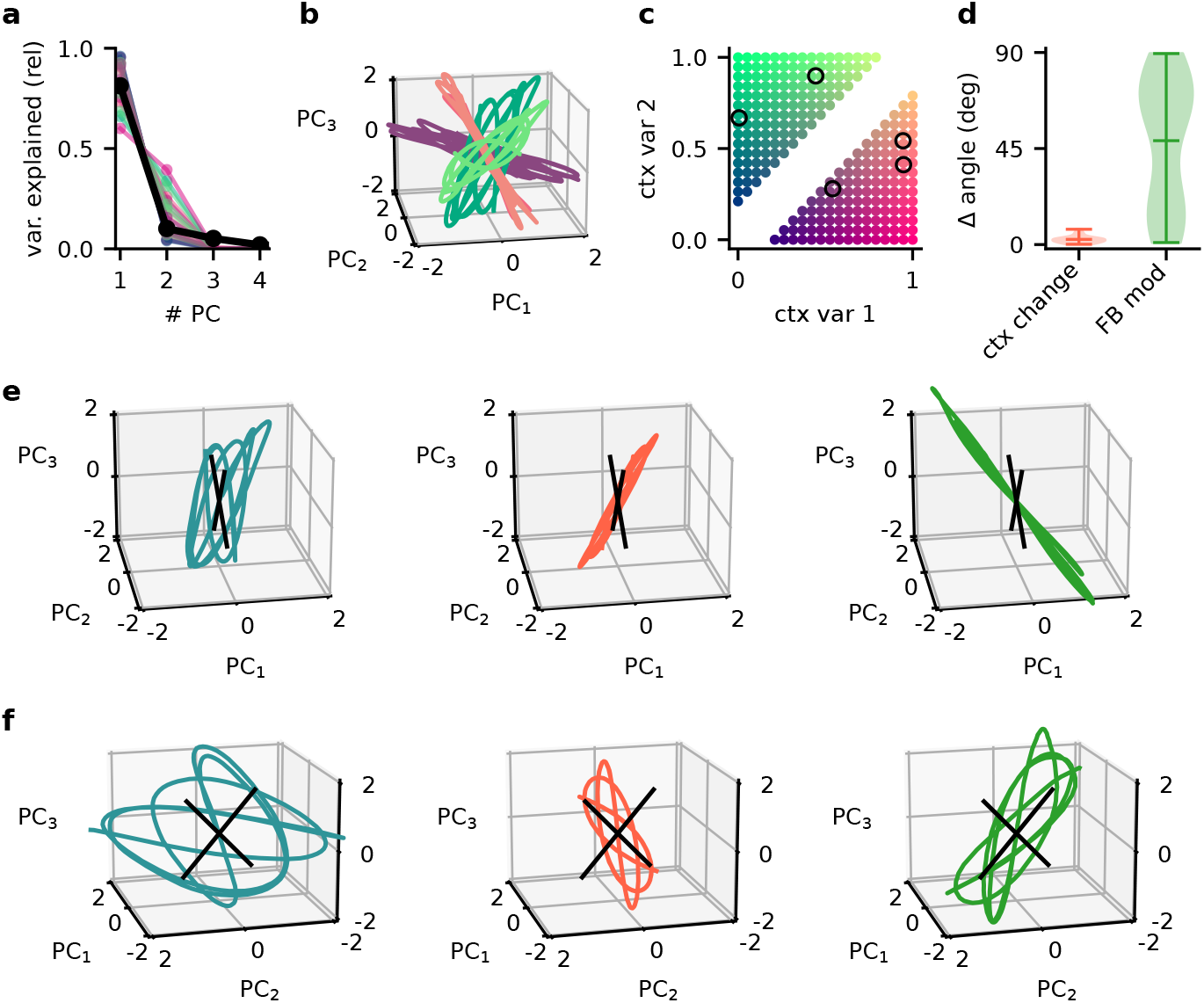
Principal component analysis captures the low-dimensional population subspaces and the subspace re-orientation with feedback. **a.** Fraction of variance explained by principal component analyses on single contexts (coloured lines) and across all contexts (black line). **b.** Population activity in the space of the first 3 PCs for 5 contexts. Colour indicates the location of the contexts in context space as shown in (**c**). **d.** Violin plot of the angle change between original subspace and the subspace for context changes (ctx change) and feedback modulation (FB mod). **e.** Population activity in the space of the first 3 PCs in different stages of the experiment. Left: context 1 with intact feedback, center: context 2 with frozen feedback, right: context 2 with intact feedback. Black lines indicate the readout vectors. **f.** Same as (e) but from a different viewpoint to show the readout space.

**Figure S10.**
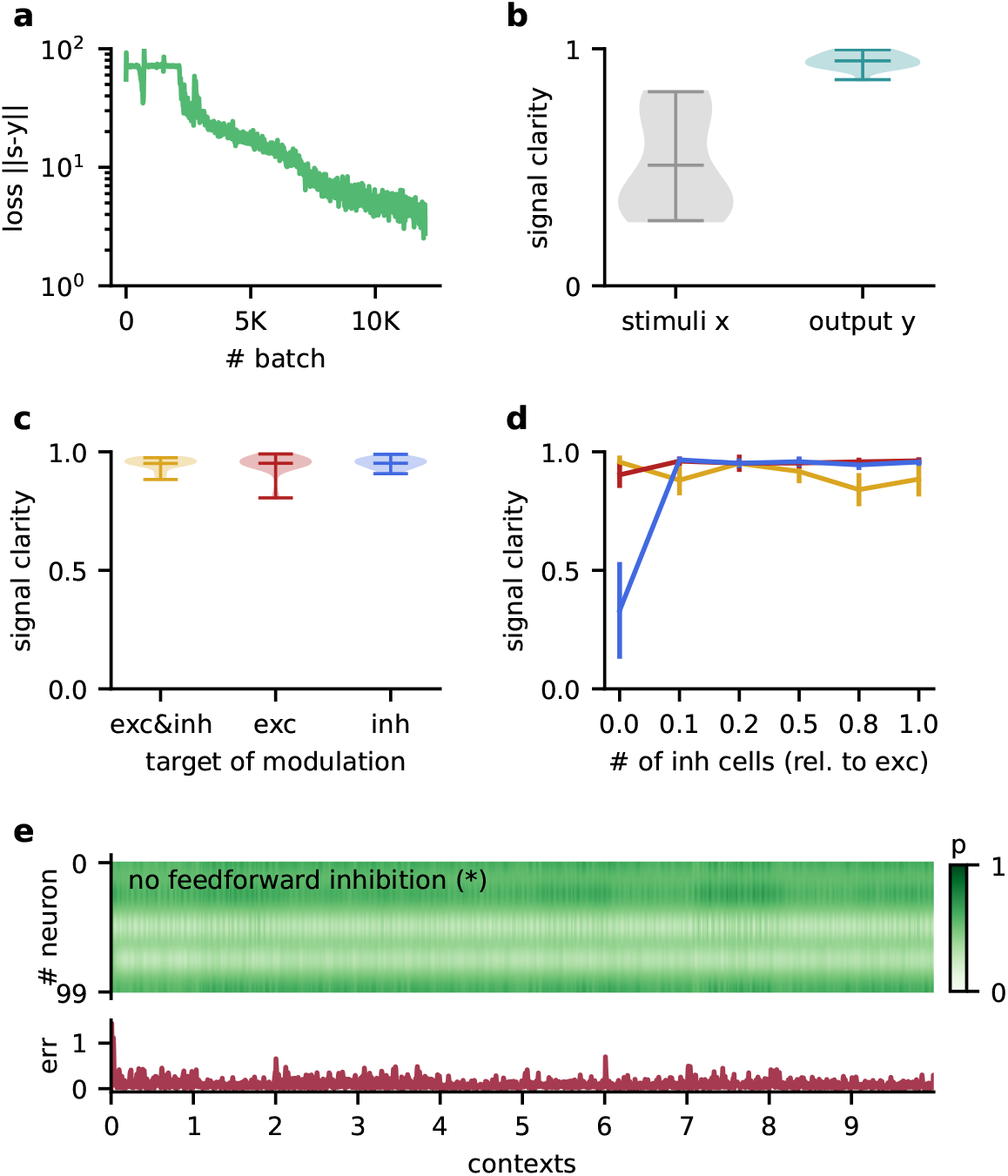
The Dalean network can learn to solve the dynamic blind source separation task and the performance of the rate-based hierarchical network does not depend on specifics of the model architecture. **a.** Loss over training. **b.** Violin plot of the signal clarity for 20 test contexts measured in the sensory stimuli and the network output. **c.** Violin plot of signal clarity for models in which excitatory, inhibitory or both types of synapses are modulated by feedback; measured over 20 contexts. **d.** Mean signal clarity across 20 contexts for different numbers of inhibitory neurons *N_I_* (relative to the number of neurons in the higher-level population). Colours correspond to the targets of modulation from (c). Error bars indicate standard deviation. The yellow arrow indicates the default parameter used in the main results. The star indicates networks without feedforward inhibition (see (e)). **e.** Top: Modulation of neurons in the higher-level population across 10 contexts without feedforward inhibition. The modulation does not switch with the context but fluctuates on a faster timescale. Bottom: Corresponding deviation of the network output from the sources.

## Notes

### Competing Interest Statement

The authors have declared no competing interest.

### Summary of Updates

The analysis of how feedback establishes the invariant subspace is now performed on a simpler network model (Fig. 4 & 5) and then validated for the Dalean network (Fig. 6). A quantification of the subspace transformation with context and feedback has been added to Fig. 5 and 6. Some results from the former Fig. 4 have been removed, to focus more on the main messages of the manuscript. The results on context decoding have been moved to a separate Fig. 7 at the end of the paper. There have been some minor changes in Fig. 1-3 (schematics and axis labels). The text has been adapted to the new structure of the manuscript. Finally, two new supplementary figures and one methods figure have been added to better illustrate how networks are trained and what they need to solve the dynamic blind source separation task.

